# Effect of heritable symbionts on maternally-derived embryo transcripts

**DOI:** 10.1101/607317

**Authors:** Mariana Mateos, Nadisha O. Silva, Paulino Ramirez, Victor M. Higareda-Alvear, Rodolfo Aramayo, James W. Erickson

## Abstract

Maternally-transmitted endosymbiotic bacteria are ubiquitous in insects. Among other influential phenotypes, many heritable symbionts of arthropods are notorious for manipulating host reproduction through one of four reproductive syndromes, which are generally exerted during early developmental stages of the host: male feminization; parthenogenesis induction; male killing; and cytoplasmic incompatibility (CI). Major advances have been achieved in understanding mechanisms and identifying symbiont factors involved in reproductive manipulation, particularly male killing and cytoplasmic incompatibility. Nonetheless, whether cytoplasmically-transmitted bacteria influence the maternally-loaded components of the egg or early embryo has not been examined. In the present study, we investigated whether heritable endosymbionts that cause different reproductive phenotypes in *Drosophila melanogaster* influence the mRNA transcriptome of early embryos. We used mRNA-seq to evaluate differential expression in *Drosophila* embryos lacking endosymbionts (control) to those harbouring the male-killing *Spiroplasma poulsonii* strain MSRO-Br, the CI-inducing *Wolbachia* strain *w*Mel, or *Spiroplasma poulsonii* strain Hyd1; a strain that lacks a reproductive phenotype and is naturally associated with *Drosophila hydei*. We found no consistent evidence of influence of symbiont on mRNA composition of early embryos, suggesting that the reproductive manipulation mechanism does not involve alteration of maternally-loaded transcripts. In addition, we capitalized on several available mRNA-seq datasets derived from *Spiroplasma*-infected *Drosophila melanogaster* embryos, to search for signals of depurination of rRNA, consistent with the activity of Ribosome Inactivating Proteins (RIPs) encoded by *Spiroplasma poulsonii*. We found small but statistically significant signals of depurination of *Drosophila* rRNA in the *Spiroplasma* treatments (both strains), but not in the symbiont-free control or *Wolbachia* treatment, consistent with the action of RIPs. The depurination signal was slightly stronger in the treatment with the male-killing strain. This result supports a recent report that RIP-induced damage contributes to male embryo death.

## 1. Introduction

Heritable associations between arthropods and endosymbiotic bacteria are widespread and influential to their hosts ^1^ and communities ^2^. With few exceptions ^3^, inheritance is achieved via the mother, generating an asymmetry in the interests of the symbiont regarding the host’s sex, whereby males are generally dead-ends for the symbiont. Consistent with this inequality, numerous heritable bacteria manipulate host reproduction in favour of symbiont-bearing females. Four such reproductive phenotypes have been described ^4^. Feminization occurs when genetic males develop and function as females. Parthenogenesis-induction occurs in haplo-diploid systems, where unfertilized eggs, which would otherwise develop as males, develop instead into reproductively functional females. In male-killing (or son-killing), the symbiont eliminates infected males to the presumed advantage of surviving infected female siblings ^5^. In cytoplasmic incompatibility (CI) ^6^, matings between symbiont-infected males and uninfected females result in death of offspring at the embryonic stage. The CI mechanism involves symbiont-mediated damage to the male sperm that is rescued in the presence of a compatible symbiont strain in the egg ^4^

Most maternally inherited symbionts are transmitted to a new host through the egg cytoplasm ^7^. As such, they may manipulate the female host’s reproductive system or usurp her cellular machinery in order to invade developing oocytes ^8–10^. Infection-induced changes during oogenesis may thus have effects on the composition of an egg (e.g. *Wolbachia* reduces the maternal transmission of the *gypsy* endogenous retrovirus ^11^). *Drosophila melanogaster* eggs contain maternal RNAs that are exclusively expressed during early development (prior to embryonic stage 5 or ~ 2h after egg deposition; AED). Zygotic transcription is silent during this period and therefore maternal mRNAs play a crucial role in early embryonic development ^12^. The egg, in a sense, is a point of convergence between the existing host, symbiont, and new host, and consequently could undergo symbiont-induced changes that could lay the foundation for the occupation of the symbiont within the new host. Furthermore, it is possible that infection by a reproductive parasite could cause changes in maternally-derived components that are necessary to induce a reproductive phenotype. As hosts of two independent lineages of maternally transmitted bacteria, *Wolbachia* and *Spiroplasma* ^13^, members of the genus *Drosophila*, have emerged as a model system for heritable symbiosis e.g. ^14,15–23^.

The wall-less bacterial genus *Spiroplasma* (class Mollicutes) is generally associated with arthropods and plants, and can reside intra- and extra-cellularly ^24^. The nature of *Spiroplasma*-host associations ranges from pathogenic to mutualistic, although the fitness consequences of the majority of *Spiroplasma* strains remain unknown. Most strains of *Spiroplasma* known to date appear to transmit horizontally from the environment or via a vector (e.g. several insect-vectored plant pathogens). A few strains of *Spiroplasma*, however, are maternally-transmitted by their arthropod hosts. Among these, ~20 species in the genus *Drosophila* are reported to harbour *Spiroplasma* ^13,25,26^. A subset of maternally-transmitted *Spiroplasma* of *Drosophila* and other arthropods are male killers. All of the *Drosophila*-associated *Spiroplasma* male-killing strains that have been genetically characterized to date fall within the *poulsonii* clade; one of the four *Spiroplasma* clades that independently invaded the genus *Drosophila* ^26^. The *poulsonii* clade also contains non-male-killing strains such as “Hyd1” and “*s*Neo”; naturally-occurring defensive mutualists of *Drosophila hydei* and *Drosophila neotestacea*, respectively ^25,27^, as well as several strains that lost the ability to kill males in the lab ^15,28,29^.

The mechanism by which *Spiroplasma* exerts death of *Drosophila* male embryos is not fully understood, but several aspects have been elucidated ^15,30–35^. First, a functional dosage compensation complex (DCC; also known as the male-specific lethal complex) is required, as mutants of components of this complex that are infected with male-killing *Spiroplasma*, fail to express male-killing ^32^. Secondly, the DCC, which normally acetylates X chromatin in males, mis-localizes to other regions of the nucleus immediately prior to the killing stage ^35^. This phenomenon is accompanied by inappropriate histone acetylation, and genome-wide transcription misregulation ^35^. Thirdly, abnormal massive apoptosis and neural disorganization occurs ^30,31,34^. The male X chromosome exhibits signs of DNA damage (chromatin bridges and segregation defects), and bridge breakage triggers sex-specific abnormal apoptosis via p53-dependent pathways ^30^. Recently Harumoto and Lemaitre ^15^ identified *spaid*, a *Spiroplasma*-encoded gene that appears to be responsible for male killing. Overexpression of *spaid* in *D. melanogaster* embryos causes death of males, but not females, and induces massive apoptosis and neural defects, reminiscent of the *Spiroplasma*-induced male-killing phenomenon in pattern, but in a somewhat delayed fashion (i.e., developmental arrest at embryonic stage 12–13 ^31,33^ in *Spiroplasma*-infected wild-type *D. melanogaster* vs. at the second larval instar with expression of the *spaid* transgene in *Spiroplasma-*free *D. melanogaster* ^15^). Spaid contains an OTU (ovarian tumor) deubiquitinase domain and ankyrin repeats (ANK). Harumoto and Lemaitre ^15^ propose that the OTU domain promotes nuclear localization of Spaid (in both female and male embryos), while the ankyrin repeats interact with DCC complex itself or with its associated histone modifications. How this leads to DNA damage and segregation defects of the male X chromosome, as well as to other phenotypes associated with male-killing described above, and which are the host cellular targets of Spaid remain unknown.

In addition to Spaid, a different type of toxin has been reported in *Spiroplasma*. Ribosome inactivating proteins (RIPs) are plant- (e.g. ricin and saporin) and bacteria-encoded (e.g. Shiga toxin) enzymes that cleave an adenine base (hereafter “depurinate”) from a specific position within a motif of the 28S rRNA that is universally conserved across eukaryotes. This motif is known as the sarcin-ricin loop (SRL). Depurination of this site renders the ribosome incapable of protein synthesis ^36^. Several confirmed and predicted RIPs are encoded in the genomes of *Spiroplasma* strains that associate with *Drosophila*, including the male-killing strain native to *D. melanogaster* (MSRO) and the closely-related non-male-killing strains *s*Neo ^17,37,38^ and Hyd1 ^39^. These *poulsonii*-clade strains are also known for their defensive abilities against certain parasitic wasps and nematodes of *Drosophila* ^21,22,27,40,41^. Evidence of SRL depurination consistent with the action of RIP has been detected in nematodes and wasps exposed to *Drosophila* harbouring *Spiroplasma* ^17,37^, and has led to the hypothesis that RIPs play a major role in the *Spiroplasma*-mediated defence against wasps and nematodes. Depurination of *Drosophila* ribosomes in *Spiroplasma*-infected treatments had been detected in larvae, but was mostly restricted to “cell-free hemolymph”, suggesting only ribosomes outside the cell were targets ^37^. Furthermore, despite detecting significant levels of depurinated *Drosophila* ribosomes in *Spiroplasma*-infected flies, this phenomenon is generally not accompanied by a significant decrease in intact (i.e., non-depurinated) ribosomes ^17,37^. This, along with the observation that the detected levels of depurination were not associated with larva-to-adult fly mortality, was interpreted as RIP activity having a negligible direct effect on fly fitness ^17,37^. A more recent study, however, revealed that *Spiroplasma*-mediated depurination of *Drosophila* ribosomes varies widely by life stage. It is strongest in embryos and old adults ^38^, but also not accompanied by a detectable decrease in intact ribosomes. In addition, significant ribosome depurination (but not significant depletion of intact ribosomes) occurs under heterologous expression of two *Spiroplasma* RIP genes in *D. melanogaster*, confirming their RIP activity. Their expression was also associated with embryo mortality (male mortality was higher), and with a reduction in fly lifespan and in adult hemocyte number ^38^.

In this study we examined whether *Spiroplasma* influences the composition of maternally loaded transcripts of host eggs (i.e., prior to fertilization), particularly in a manner that may facilitate male-killing. To do so we compared the effects of three heritable endosymbionts (a male-killing *Spiroplasma*, a non-male-killing *Spiroplasma*, and a CI-inducing *Wolbachia*) on the composition of mRNA transcripts of early embryos of *Drosophila melanogaster*. Furthermore, we used data from this study and published RNA-seq data to analyse whether there is a difference in signals of rRNA depurination of host embryo ribosomes in the presence of male-killing and non-male-killing symbionts.

## 2. Materials and Methods

### 2.1 Generation of symbiont treatments

We used three infection treatments, one *Wolbachia* and two *Spiroplasma* strains, and a symbiont-free control (Fig. 1). Laboratory stocks of *D. melanogaster* (Canton S strain; CS) that naturally harbour the *Wolbachia* strain *w*Mel were used to generate the *Wolbachia* treatment (W+S−). Positive infection for *w*Mel was confirmed based on PCR with *Wolbachia*-specific primers targeting the *wsp* gene ^42^. The same stock was reared in tetracycline food (final concentration 0.02g/ml) for two generations, followed by three generations of antibiotic-free food to generate a *Wolbachia*-free (W−) stock. The W− flies served as the symbiont-free control. The *Spiroplasma* infection treatments (W−S+) were generated by artificially infecting *Wolbachia*-free (W−) flies with the strain MSRO-BR (Red 42) native to *D. melanogaster* ^43^ or Hyd1 (TEN-104-106) native to *D. hydei* ^13^. Fifteen *Wolbachia*-free (W−) females (15 lines) were infected per *Spiroplasma* strain. These artificially infected lines were maintained for 3–5 generations before being used for the experiment. *Spiroplasma*-infected (W−S+) lines were selected every generation to ensure positive infection status, based on PCR with *Spiroplasma*-specific 16S ribosomal DNA primers ^43^. MSRO treatment lines were backcrossed to *Wolbachia*-free (W−) CS males every generation, as male-killing by this strain is nearly perfect. A minimum of four infected lines was combined, per replicate, at the start of the experiment, to create a total of three biological replicates for each *Spiroplasma* treatment. The biological replicates for the *Wolbachia* treatment and control were maintained as three different populations for four generations prior to the start of the experiment.

**Figure 1.**
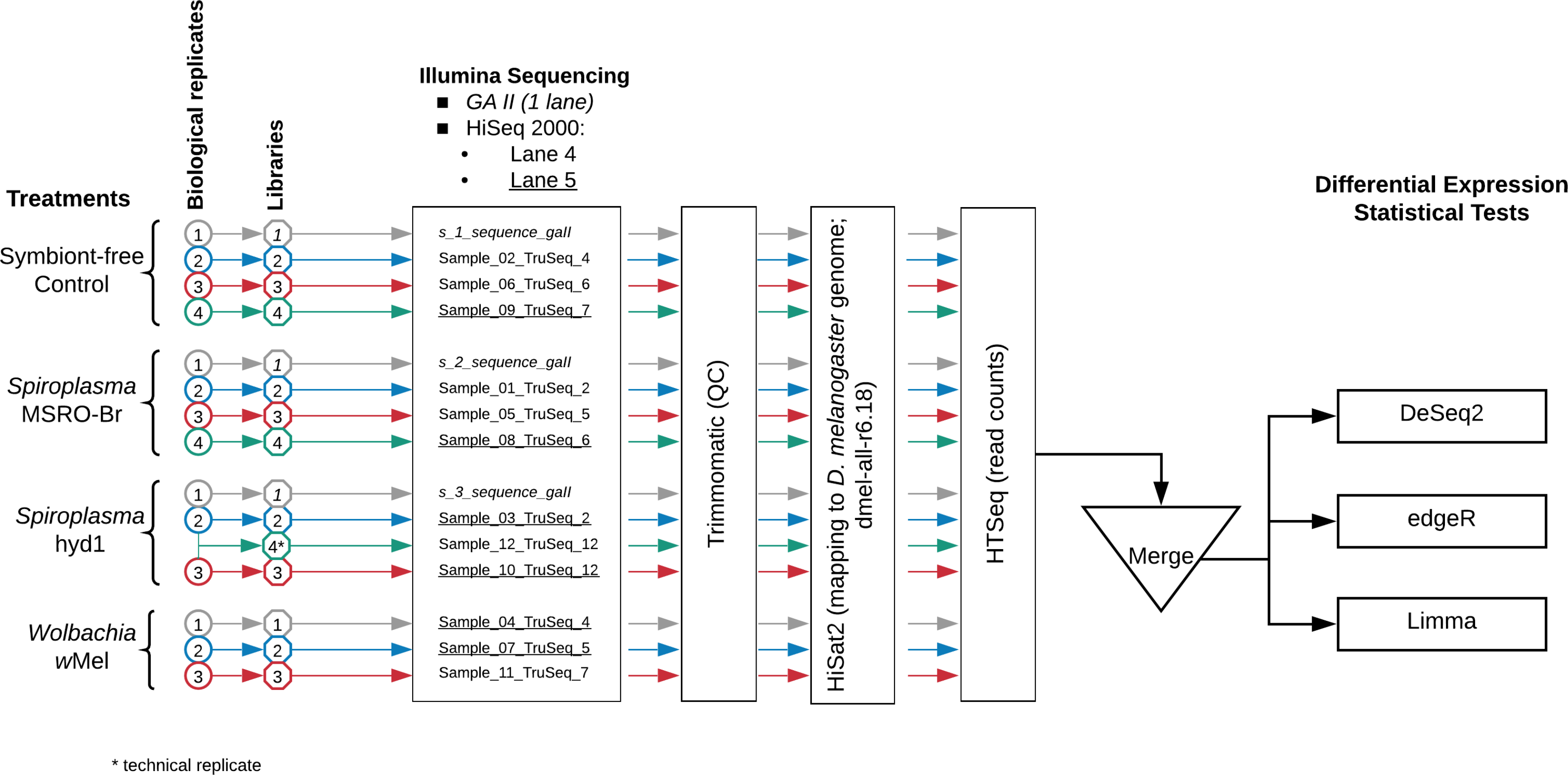
Experimental design and workflow for data analysis. The biological replicates, corresponding libraries, and sequence file labels are distinguished by colours and font type. Italics = samples run on Illumina GAII. Non-italics = samples run on Illumina HiSeq 2000; non-underlined samples were pooled into one sequencing lane (4), whereas underlined samples were pooled into another lane (5). Library labelled “Sample_12_TruSeq_12” is the result of combining total RNA from Hyd1 biological replicates 2 and 3, and thus considered a technical, rather than biological, replicate. This library was excluded from the differential expression analyses.

### 2.2 Embryo collection

Approximately 40–50 three-day-old virgin females, from each replicate of each treatment, were allowed to mate in cages with *Wolbachia*-free (W−) CS males during the collection period, and allowed to lay eggs on cornmeal food plates. The initial batch of eggs was discarded to improve the chances of collecting fertilized eggs from the same stage ^44^. Thereafter, egg laying was monitored and cornmeal plates were changed approximately every 45 min, so as to collect embryos that were on average ~60–75 min old. Eggs were collected from each replicate with a small brush 6–8 times over a 2-day period for each treatment and the control. The eggs were placed in sterile RNase-free 1.7 ml microtubes, and immediately put on dry ice during the collection period, after which they were transferred to −80°C for storage.

### 2.3 RNA extraction, library preparation, and sequencing

Three to four biological replicates per condition (MSRO, Hyd1, wMel, and control) were used for the extractions (see Fig. 1). RNA was extracted per collection tube of eggs (mentioned above) with Trizol® Plus RNA Purification System (Invitrogen, Carlsbad, CA) according to the manufacturer’s protocol. RNA from tubes that belonged to the same biological replicate within a treatment was pooled. All RNA samples were DNase-treated with Ambion® DNA- free (Invitrogen) to remove any DNA contamination. Total RNA was quantified with a NanoDrop® ND-1000 UV spectrophotometer (NanoDrop Technologies, Wilmington, DE), and sample quality and integrity were further tested with the Agilent 2100 BioAnalyzer (Agilent Inc., Santa Clara, CA).

Total RNA was submitted to the Texas AgriLife Genomics and Bioinfomatics Services facility for library preparation, sequencing, and demultiplexing. Three biological replicates were subjected to multiplex NuGen library preparation and pooled into a single sequencing lane of the Illumina (San Diego, CA) Genome Analyzer II platform (single-end; 76 bp read length; italicized samples in Fig. 1). The remaining biological replicates were subjected to the Illumina mRNA TruSeq kit library preparation protocol (12 libraries). Half of the libraries were pooled into one lane of the Illumina HiSeq 2000 platform and the other half were pooled into another lane (single-end; 100 bp read length; see Fig. 1).

### 2.4 Analyses of Differential Expression (DE)

Raw reads have been deposited in the NCBI SRA Database under Accession Numbers SRR7279355–SRR7279369 (BioProject PRJNA474708; BioSample SAMN09370647). Command lines, as well as input and output files needed to replicate our analyses, from the different pipelines used are available in Supporting Data Files S1–S4. Raw RNA sequence files were first processed with Trimmomatic (v 0.35) ^45^ to remove adapters and low quality reads (see Fig. 1). The processed runs were then mapped to the Flybase *Drosophila* genome (version dmel-all-r6.18) with HiSat2 (v.2.0.5) ^46^ to obtain bam files. The files containing the mapped read information were then analysed with HTseq (0.6.1) ^47^ to obtain read counts for both genes and exon regions specified in a corresponding gff file for the dmel-all-r6.18 genome sequence. The resulting count files were then manually formatted into a count matrix suitable for differential expression analysis (Supporting Data File S2).

Differential expression analyses were performed with three different pipelines edgeR v.3.20.5 ^48^ and Limma+Voom v. limma_3.34.5 ^49^ and DeSeq2 v.1.18.1 ^50^. Command line and relevant input files for these analyses are provided in Supporting Data Files S1–S3. All programs conducted their respective tests on genes/exons with 10 or more mapped reads per replicate within at least one of two the treatments being compared. For the EdgeR pipeline both the glm and QFglm models were used for DE. For Deseq2, the wald statistical test was used for DE analysis. Before analysis in both Limma-Voom and EdgeR pipeline, two genes with extremely high counts, 16S rRNA (FBgn0013686) and ef-1α (FBgn0284245), were removed. Genes with an adjusted *p*-value (or false discovery rate; FDR) of 0.05 or less were considered to be potentially differentially expressed (DE).

### 2.5 Power Analyses for Differential Expression

To identify possible limitations of our experimental design in the detection of DE genes, we performed a statistical power analysis with the method of Hart et al. ^51^. The parameters (and rationale) used to run the simulations are provided in Table 1 and Fig. 2.

**Table 1.**
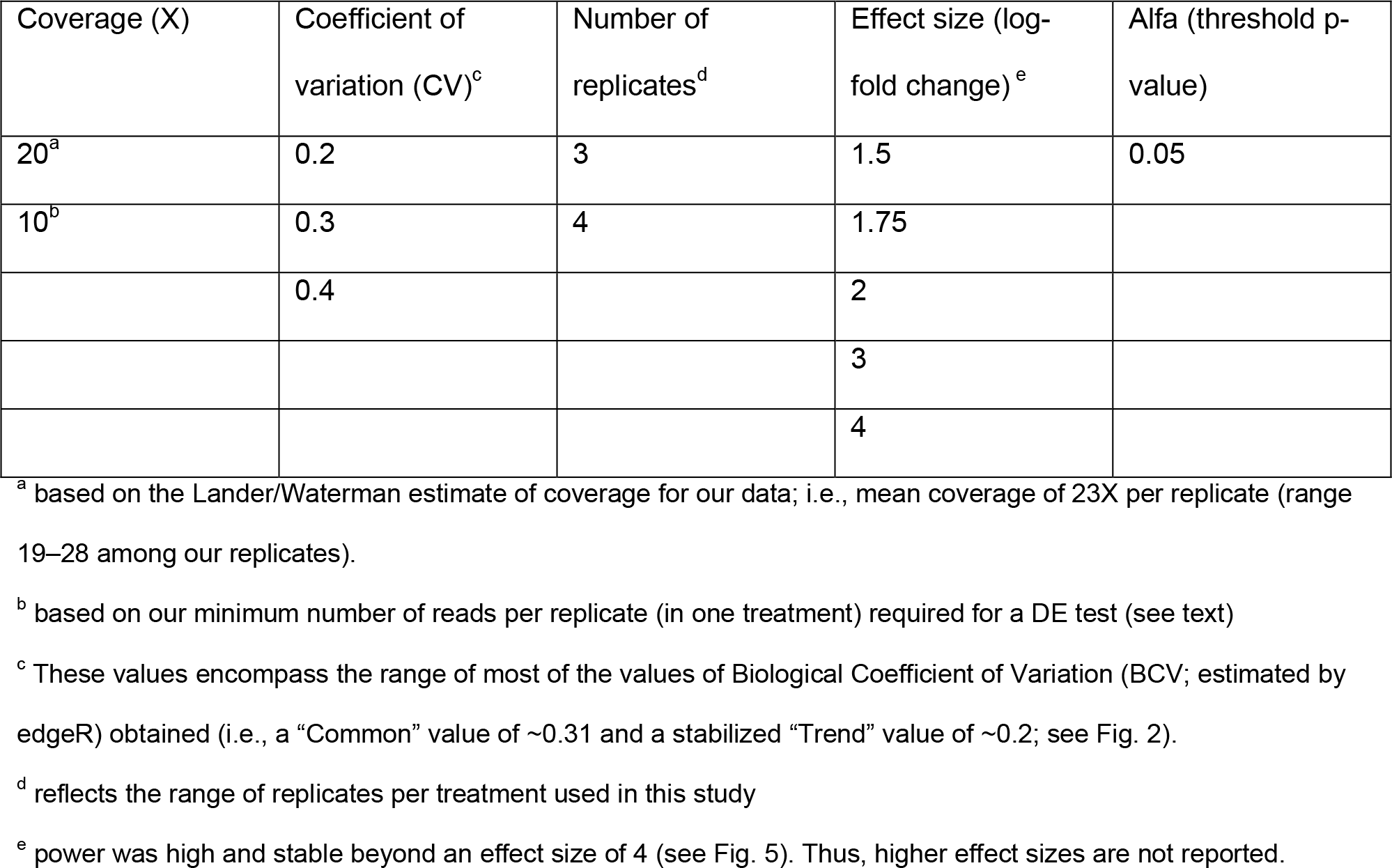
Parameters used to run the simulations of the power analyses. All possible combinations of these parameters (60 total) were run.

| Coverage (X) | Coefficient of variation (CV)<sup>c</sup> | Number of replicates<sup>d</sup> | Effect size (log-fold change)<sup>e</sup> | Alfa (threshold p-value) |
| --- | --- | --- | --- | --- |
| 20<sup>a</sup> | 0.2 | 3 | 1.5 | 0.05 |
| 10<sup>b</sup> | 0.3 | 4 | 1.75 |  |
|  | 0.4 |  | 2 |  |
|  |  |  | 3 |  |
|  |  |  | 4 |  |

<sup>a</sup> based on the Lander/Waterman estimate of coverage for our data; i.e., mean coverage of 23X per replicate (range 19–28 among our replicates).
<sup>b</sup> based on our minimum number of reads per replicate (in one treatment) required for a DE test (see text)
<sup>c</sup> These values encompass the range of most of the values of Biological Coefficient of Variation (BCV; estimated by edgeR) obtained (i.e., a “Common” value of ~0.31 and a stabilized “Trend” value of ~0.2; see Fig. 2).
<sup>d</sup> reflects the range of replicates per treatment used in this study
<sup>e</sup> power was high and stable beyond an effect size of 4 (see Fig. 5). Thus, higher effect sizes are not reported.

**Figure 2.**
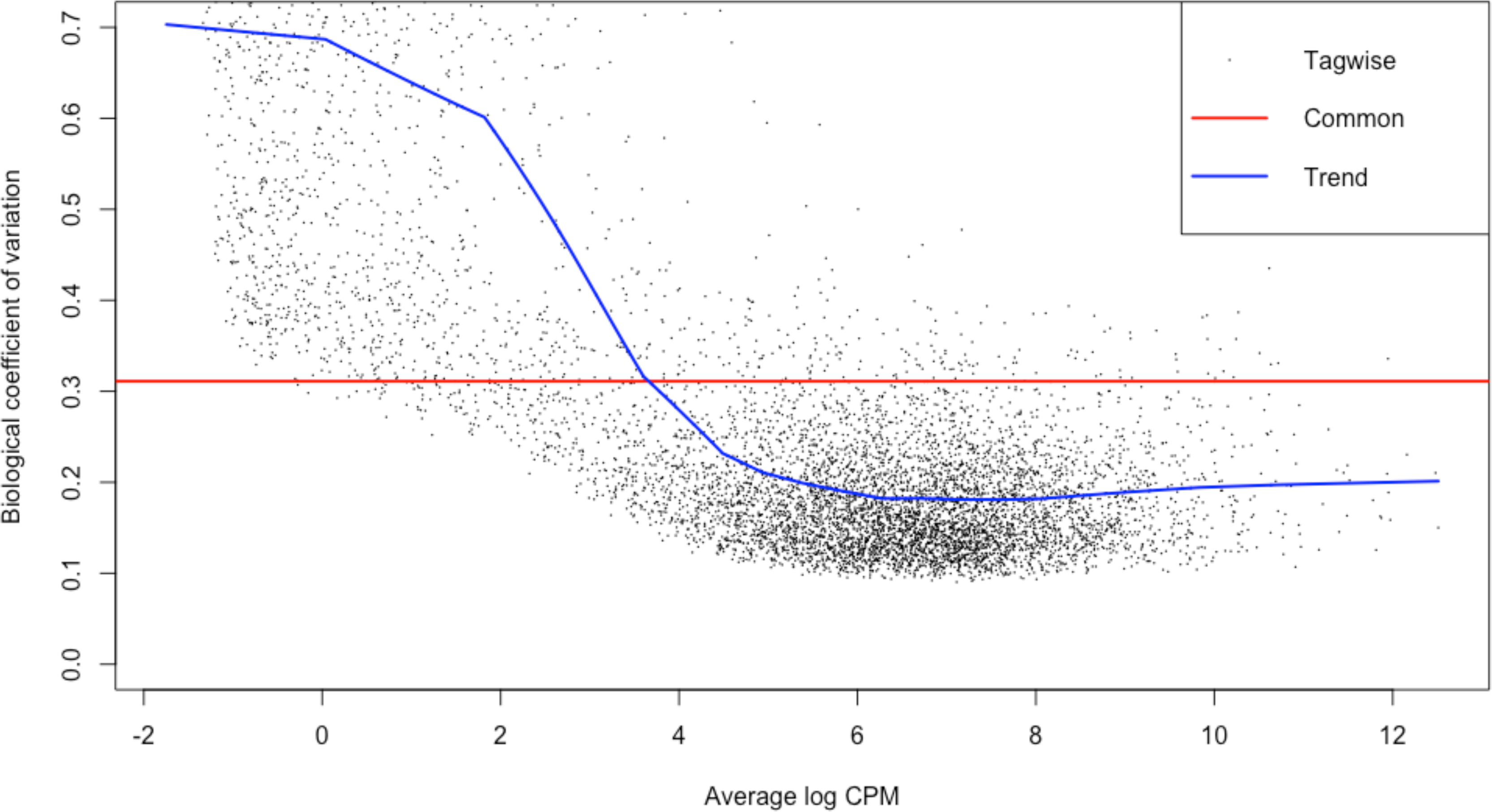
Biological coefficient of variation (BCV) for our data plotted against average log CPM (copies per million), as estimated by edgeR.

### 2.6 Analyses of depurination signal in the sarcin-ricin loop (SRL) of the 28S rRNA of *Drosophila*

RIP toxins remove a specific adenine present in the sarcin-ricin loop (SRL) of the 28S rRNA leaving an abasic site (i.e., the backbone remains intact) ^52^. When a reverse transcriptase encounters an abasic site, it preferentially adds an adenine in the nascent complementary (c)DNA strand ^53^. This property, which results in an incorrect base at the RIP-depurinated site in the cDNA and all subsequent PCR amplification steps, can be used to detect evidence of RIP activity in any procedure that relies on reverse transcription (e.g. RNA-seq or reverse-transcription qPCR). To examine whether a signal of depurination consistent with RIP activity was detectable in *Spiroplasma*-infected flies, we used the RNA-seq data generated in the present study, as well as RNA-seq data (also derived from poly-A-tail-enriched RNA libraries) from two published studies that also compared patterns of gene expression from *Spiroplasma*-infected and *Spiroplasma*-free *D. melanogaster* embryos: NCBI Acc. No. PRJDB4469 ^30^; and PRJNA318373 ^35^. The *Spiroplasma* strain examined by ^35^ and ^30^ is substrain UG (for Uganda) of MSRO. Bowtie2 v.2.1.0 ^54^ and Samtools 0.1.19 ^55^ were used to map and extract, respectively, the reads from the three studies to the 28S rRNA gene of *D. melanogaste*r (Acc. No. NR_133562.1 positions 2591–3970). We verified that the reads that mapped to this region did not also map to other parts of the *Drosophila* genome or to the genomes of *Spiroplasma* or *Wolbachia*. To visualize and count the shift from A to T (or other bases) in the source RNA pool, the extracted reads were mapped again to the same reference sequence in Geneious v.11.1.2 (Biomatters Inc., Newark, NJ; “low sensitivity mode”; maximum gap size=3; iterate up to 25 times). The number of reads containing each of the four bases or a gap at the target site was counted (gapped reads were excluded from subsequent analyses). The proportion of reads with an A at the target site (i.e., putatively intact rRNA) was calculated and compared among treatments and replicates. We used a mixed model in JMP Pro 13 (SAS Institute Inc., Cary, NC) to examine the effect of symbiont treatment (fixed) on the proportion of adenines (arcsine square root transformed) at the target position (excluding gaps from total number of reads). Source Study was treated as a random effect.

## 3. Results

The present study aimed at determining whether heritable symbionts alter the composition of maternally-loaded mRNAs in *D. melanogaster*. To achieve this, we used mRNA-seq to evaluate differential expression (DE) in *Drosophila* embryos lacking endosymbionts (control) to those harbouring one of the following heritable endosymbionts: the male-killing *Spiroplasma poulsonii* strain MSRO-Br; the CI-inducing *Wolbachia* strain *w*Mel; and the non-male-killing *Spiroplasma poulsonii* strain Hyd1 (native to *D. hydei*). A power analysis was performed to determine the limitations of our experimental design. A secondary goal of this study capitalized on several available mRNA-seq datasets derived from *Spiroplasma*-infected *Drosophila melanogaster* embryos, to search for signals of damage (i.e., depurination) to *Drosophila* rRNA, consistent with the activity of Ribosome Inactivating Proteins (RIPs), which are encoded in the genomes of several *Spiroplasma* strains. This assay examined the proportion of adenines at the RIP target position (indicative of intact rRNA) in *Spiroplasma*-infected vs. *Spiroplasma*-free treatments.

### 3.1 Differential Expression

The number of reads that was obtained, passed QC, and mapped to the *Drosophila* genome is shown in Table 2. The PCA plot did not reveal any particular grouping of samples according to treatment (Fig. 3). PC1 explained 51% of the variance and revealed a slight “batch” effect, where the “GAII” replicates (i.e., 1_gall, 2_gall, and 3_gall) fell to the left of the other replicates. Nonetheless, removal of these replicates did not lead to a better grouping of replicates within each treatment (not shown). Thus, we considered that exclusion of the “GAII” replicates was not justified. Inclusion or exclusion of the Hyd1 technical replicate (e.g. n = 4 vs. 3 replicates, respectively) did not affect the differential expression analyses (only n = 3 is shown).

**Table 2.** Number of reads obtained, retained after quality control, and mapped to the reference genome.

| Treatment | Replicate | Total reads | Reads after quality control |  | Reads mapped to reference genome |  |
| --- | --- | --- | --- | --- | --- | --- |
|  |  | Number | Number | % | Number | % |
| Control | 1_gall | 38,623,190 | 38,595,804 | 99.93 | 36,658,295 | 94.98 |
| Control | 2_Truseq_4 | 32,414,845 | 32,413,140 | 99.99 | 31,382,402 | 96.82 |
| Control | 6_Truseq_6 | 28,893,141 | 28,891,426 | 99.99 | 27,923,563 | 96.65 |
| Control | 9_Truseq_7 | 31,672,204 | 31,672,204 | 100.00 | 30,671,362 | 96.84 |
| Hyd1 | 3_gall | 39,050,068 | 39,031,083 | 99.95 | 36,931,211 | 94.62 |
| Hyd1 | 3_Truseq_2 | 28,074,997 | 28,074,997 | 100.00 | 27,151,330 | 96.71 |
| Hyd1 | 10_Truseq_12 | 31,084,062 | 31,084,062 | 100.00 | 30,070,722 | 96.74 |
| MSRO | 2_gall | 36,838,900 | 36,824,546 | 99.96 | 34,640,850 | 94.07 |
| MSRO | 1_Truseq_2 | 31,841,684 | 31,832,465 | 99.97 | 30,867,941 | 96.97 |
| MSRO | 5_Truseq_5 | 30,537,077 | 30,535,822 | 100.00 | 29,442,640 | 96.42 |
| MSRO | 8_Truseq_6 | 30,124,318 | 30,124,318 | 100.00 | 29,172,390 | 96.84 |
| wMel | 4_Truseq_4 | 32,691,282 | 32,691,282 | 100.00 | 31,449,013 | 96.20 |
| wMel | 7_Truseq_5 | 27,032,887 | 27,032,887 | 100.00 | 26,073,220 | 96.45 |
| wMel | 11_Truseq_7 | 28,550,836 | 28,548,722 | 99.99 | 27,595,195 | 96.66 |

**Figure 3.**
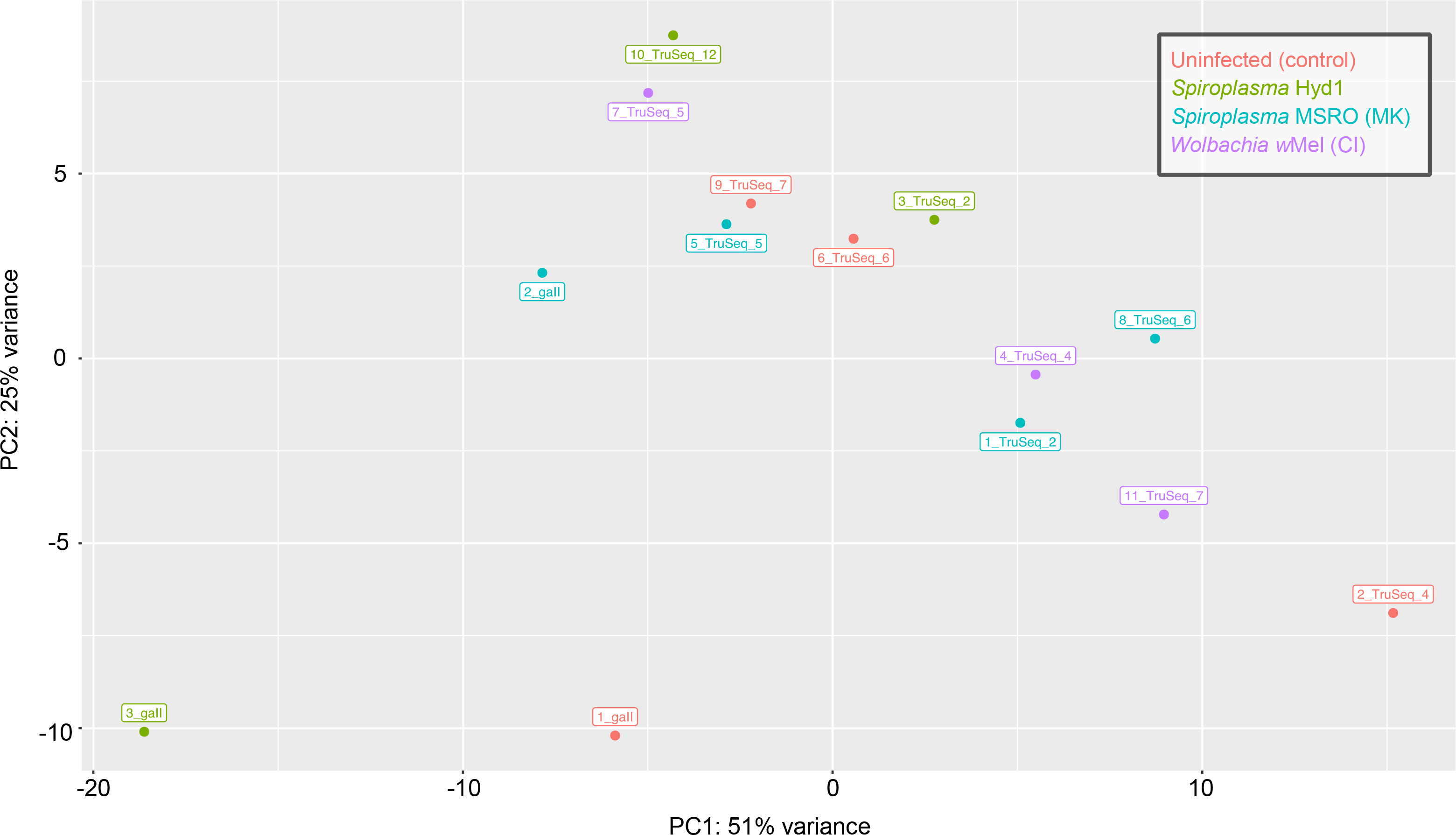
DESeq-generated principle component analysis (PCA) with VSD transformation on the 14 biological replicates (see Fig. 1). Each treatment is labelled by a different colour. MK = male killing; CI = cytoplasmic incompatibility. The technical replicate “Sample_12_TruSeq_12” (not shown) was intermediate between “Sample_3_TruSeq_2” and “Sample_10_TruSeq_12”.

All differential expression (DE) tests were performed by comparing each symbiont treatment to the symbiont-free control. None of the eight pipelines used detected differentially expressed (DE) genes in the MSRO (male-killing *Spiroplasma*) and wMel (CI-inducing *Wolbachia*) treatments (Table 3). For the Hyd1 (non-male-killing *Spiroplasma*) treatment, half of the pipelines did not detect DE genes, whereas the other half detected a few DE genes (Table 3). The Deseq2 pipeline and the edgeR pipeline (glm model only) detected 5–12 differentially expressed (DE) genes in the Hyd1 treatment based on ‘by gene’ and ‘by exon’ analyses (Table 3). Only five DE genes (*ect*, *Osi6*, *Osi7*, FBgn0038339, and FBgn0037099) were found in common among these pipelines. For all of these, the Hyd1 treatment exhibited lower expression, and lower variation among replicates, than the other treatments (Table 3 and Fig. 4).

**Table 3.**
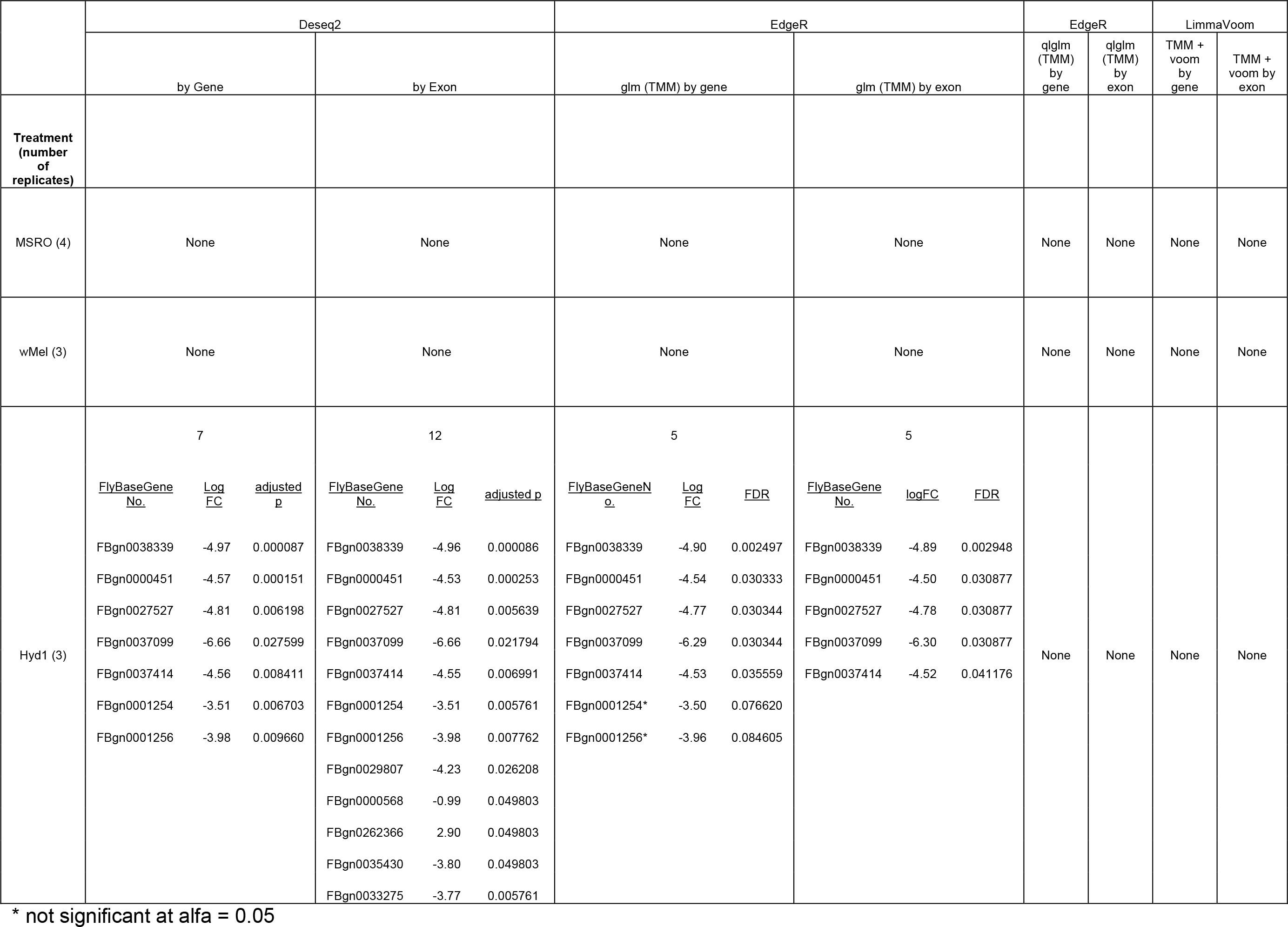
The number, identity (Flybase gene number), log fold-change (logFC), adjusted *p*-value or FDR, of genes identified as differentially expressed (DE) between each treatment and the symbiont-free control, by each of the four methods utilized. Results are shown for analyses of both genes and exons. Only the Hyd1 treatment exhibited significant DE genes.

**Figure 4.**
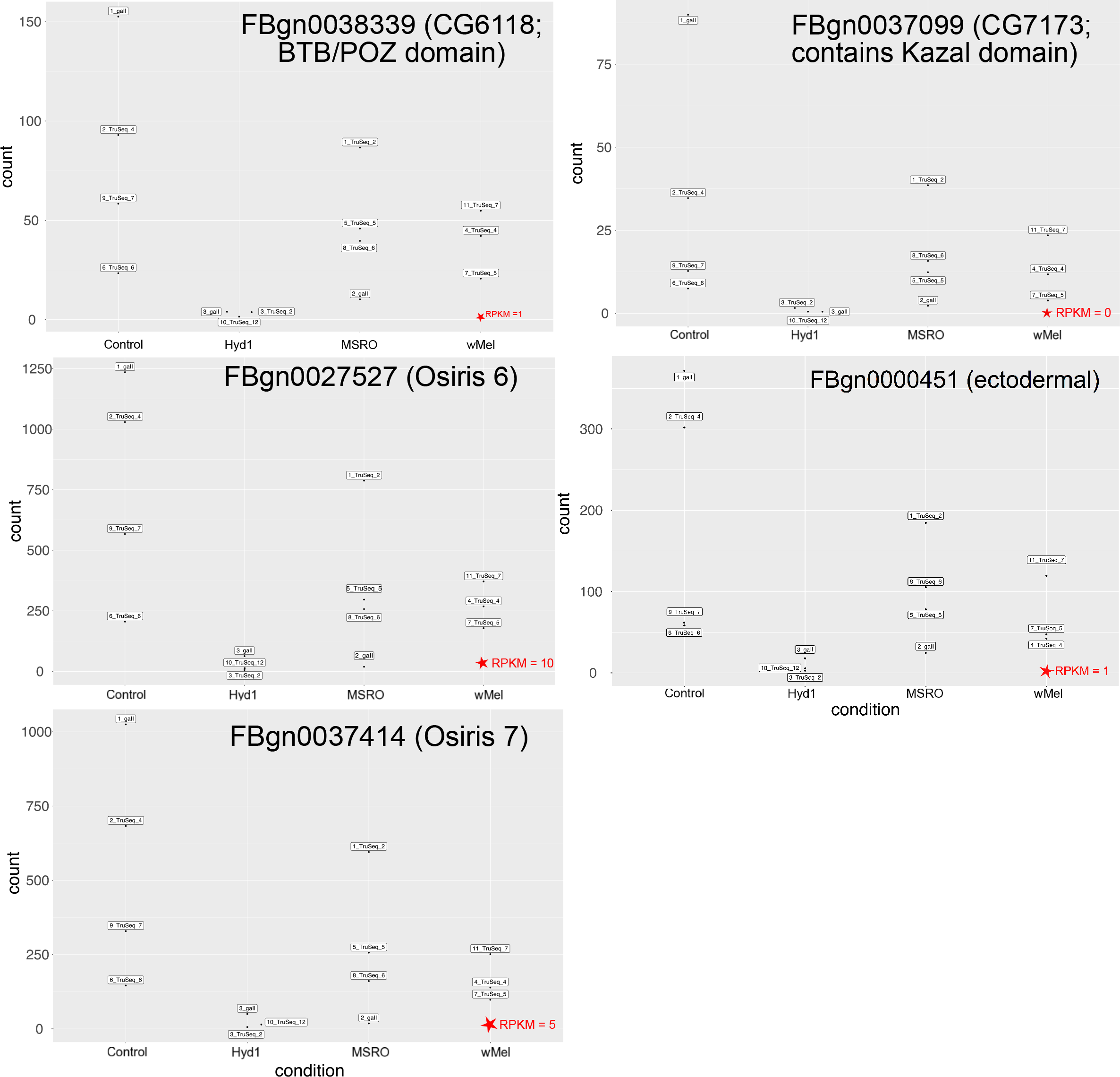
Normalized counts, as estimated by Deseq2, for the five genes detected as differentially expressed (DE) by a subset of the pipelines (see text and Table 3). Red star indicates the RPKM value reported by FlyBase for the equivalent treatment (0-2 h embryo; *Wolbachia*-infected).

### 3.2 Power Analyses for Differential Expression

A plot of the power analyses is shown in Fig. 5 and the rationale for the parameters assumed is provided in Table 1. Assuming a coverage of 20X and a coefficient of variation of 0.3, we should expect to detect: ~100% of genes differentially expressed by log-fold changes ≥ 3; at least 60% of genes with log-fold changes ≥ 2, and at least ~40% of genes with log-fold changes ≥ 1.75. Therefore, these results suggest that our design and data have sufficient power to detect log-fold changes ≥ 3, but more limited power below that log-fold change value.

**Figure 5.**
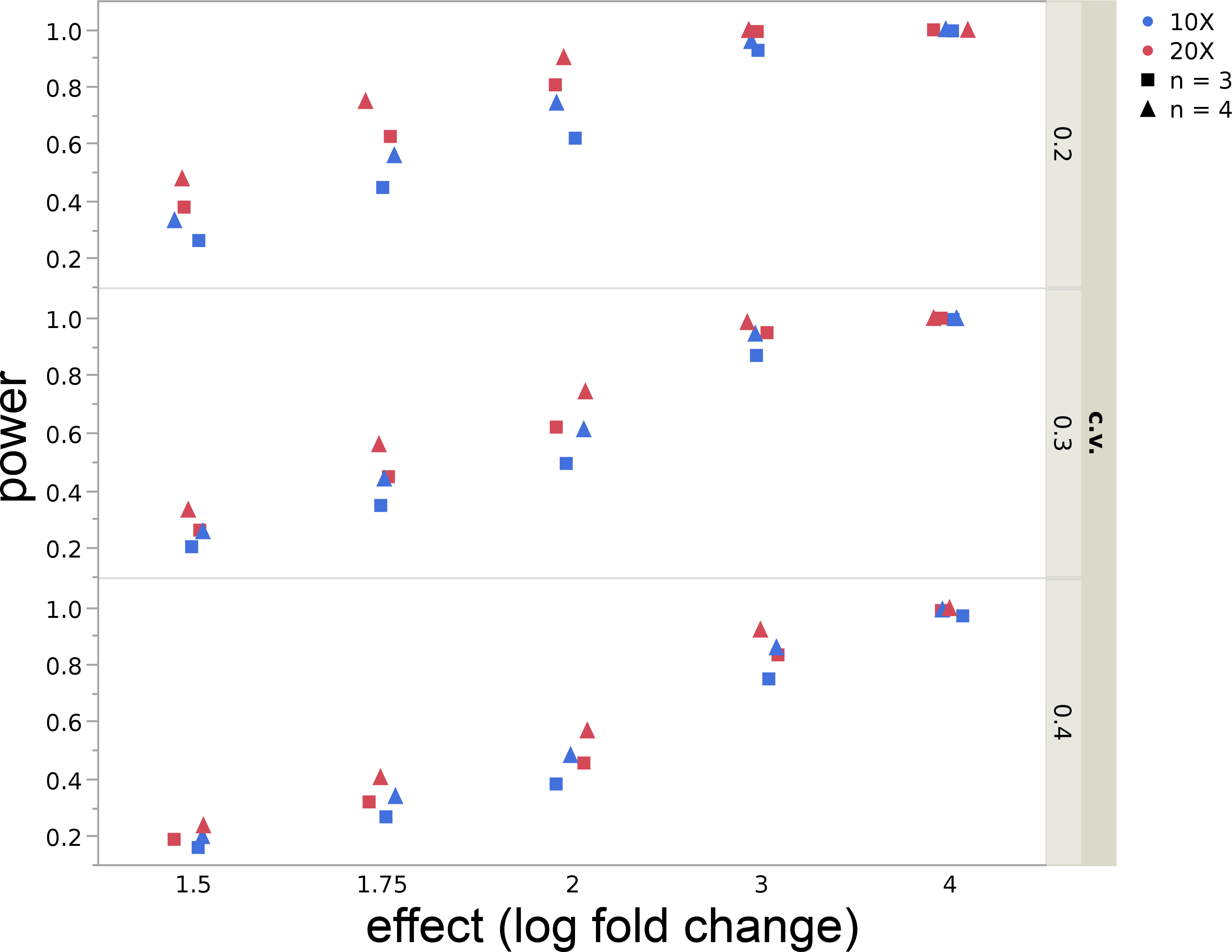
Power Analyses. Estimation of power for alfa = 0.05, at five different effect values (= log-fold change: 1.5–4); number of replicates (n = 3 and 4), coverage values (X = 10 and 20), and coefficient of variation (c.v. = 0.2, 0.3, 0.4).

### 3.3. Signals of Depurination

The number of reads mapped to the target site, as well as the proportion of adenines (representative of non-depurinated rRNA) by treatment replicate, are shown Fig. 6 and Table S1. Symbiont treatment had a significant effect on the proportion of adenines (*F*_4,26_ = 29.8; *p* < 0.0001), whereas source study did not (Wald *p* = 0.4895). Both MSRO (UG and BR) treatments had a significantly (*p* ≤ 0.001) smaller proportion of As (mean = 96.78%) than the control (mean = 99.79%) and *w*Mel (mean = 100%) treatments. Hyd1 exhibited intermediate proportions of As (mean = 98.53%; Fig. 6A). In other words, we detected a mean depurination of 3.22% for MSRO and 1.47% for Hyd1, and effectively no depurination for the control or wMel treatment. Pooling of the two substrains of MSRO (i.e., MSRO-UG and MSRO-BR) into one treatment produced similar results (not shown). The Harumoto et al. ^30^ dataset, which included two embryo stages and differentiated between male and female embryos, did not exhibit substantially different depurination levels between stages or sex (e.g. mean %A was 98.27 for females and 98.49 for males; see Table S1).

**Figure 6.**
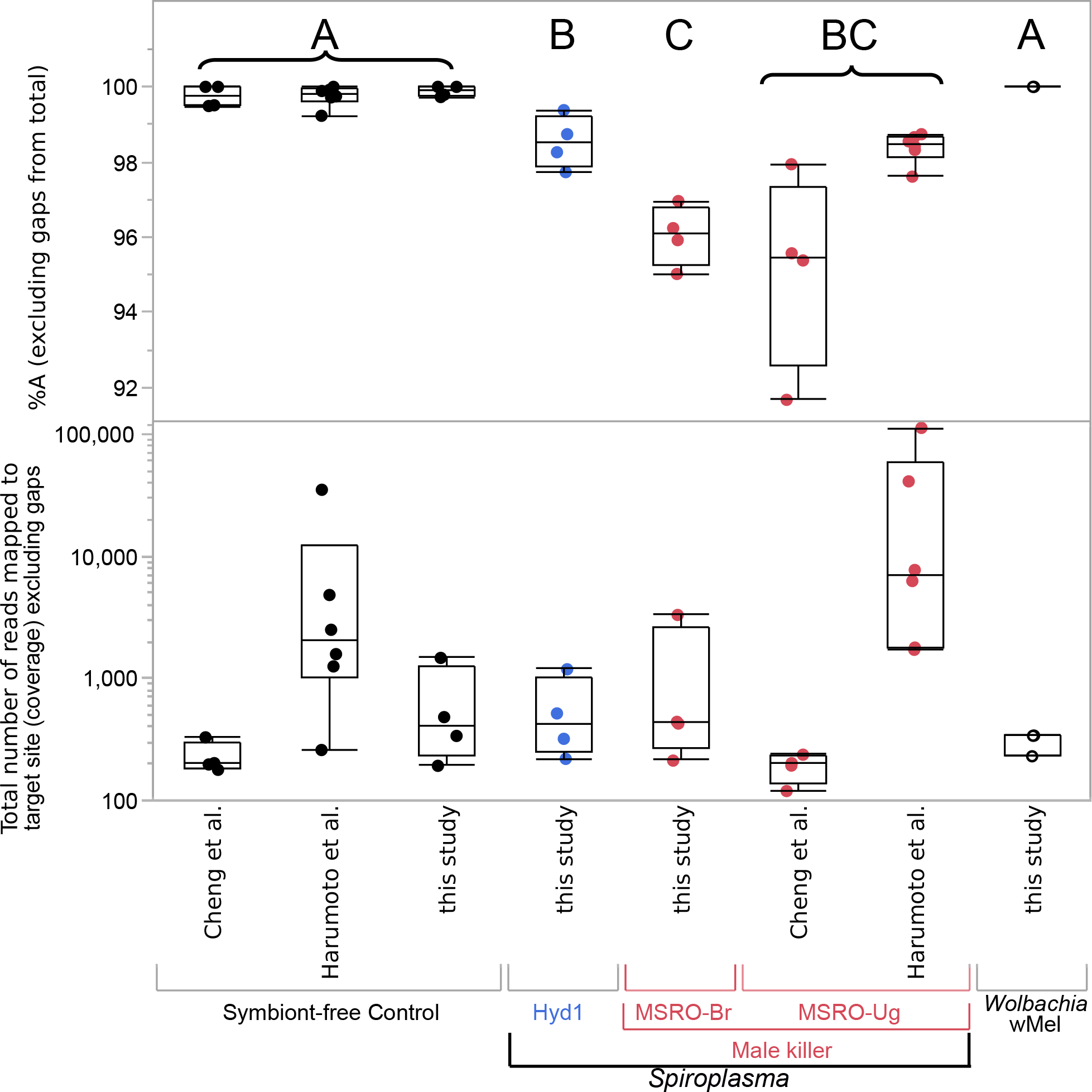
Presence of male-killing *Spiroplasma* (MSRO) results in a significant signal of depurination. Top Panel. Percent of adenines (i.e., the expected base in the absence of depurination) at the target site of Ribosome Inactivating Proteins (RIPs). Different letters indicate significant Posthoc tests (at P-value < 0.04). Bottom Panel. Coverage (number of reads) that mapped to the target depurination sites. Jitter points (replicates) and box plots per treatment (symbiont and source study) are shown.

### 3.4 Outcome of mapping mRNAseq reads to symbiont genomes

To determine the degree to which poly-A-tailed enriched RNAseq data yield reads assignable to the bacterial symbionts, we used Bowtie2 (v.2.3.4) ^54^ with default settings to map the reads from each symbiont-infected replicate to the closest symbiont reference genome available. The number of reads per replicate mapped to the symbiont genome ranged from 5 (for a Hyd1 replicate) to 8619 (for a wMel replicate; Table S2), representing a very small fraction of the total reads. The percentage of these putative symbiont-derived reads that mapped to the symbiont ribosomal genes was quite variable (range 3.94–100%; mean 52%). The above numbers of symbiont-derived genes are inadequate for analyses of differential expression (DE) of the symbiont. Collectively these findings suggest that libraries generated via a poly-A-tail enrichment process tend to yield a very low proportion of reads of bacterial origin; a result that is expected given that polyadenylation is not part of the process of bacterial mRNA maturation.

## 4. Discussion

### 4.1 Effects of symbionts on early embryo mRNA composition

The present study aimed to test whether infection by a heritable bacterium strain that kills males during the embryonic stage influences the composition of mRNA transcripts present in the early embryo (ca. 60-75 min post-oviposition), and thus, before the onset of zygotic transcription. Based on several analysis pipelines and assumptions, we did not detect any differential expression in genes among embryos harbouring the male-killing *Spiroplasma* strain (MSRO-Br), the CI-inducing *Wolbachia* strain, and the symbiont-free control. These results suggest that these symbionts do not influence the composition of initial maternally-loaded transcripts or their degradation up to the examined stage, unless they do so to a degree below what was detectable by our experimental design (e.g. below a log-fold change of 3; see Power Analyses). Furthermore, it is possible, that such cytoplasmically transmitted symbionts could influence the composition of maternally-loaded proteins, or regulate the protein complement of the early embryo in a transcriptionally-independent manner (e.g. through post-transcriptional or post-translational controls). To our knowledge, these features have not been compared in *Spiroplasma*-infected vs. uninfected embryos.

Our results showing no effect of the male-killing *Spiroplasma* on maternally-loaded mRNA composition are consistent with what is known about the male-killing mechanism. A *Spiroplasma*-encoded protein (Spaid) appears to interact with the DCC, which assembles onto the X chromosome of wild-type males but not females, and is accompanied by DNA damage and abnormal apoptosis ^15^. This model does not require “priming” of the egg during oogenesis by the symbiont. Nonetheless, the delayed expression of male death in males flies over-expressing wild type Spaid (2nd larval instar), compared to those harbouring male-killing *Spiroplasma*, requires further investigation and may indicate that other *Spiroplasma*-encoded factors are relevant to the male-killing phenotype (e.g. RIP) ^38^.

The differential expression analyses of the comparisons involving the Hyd1 treatment (i.e., the non-male-killing *Spiroplasma* strain native to *D. hydei*), yielded equivocal results, with four pipelines revealing no differentially expressed genes, and the other four pipelines revealing an intersection of five DE genes (*ect*, *Osi6*, *Osi7*, FBgn0038339, and FBgn0037099). All of these five genes had a lower expression level in the Hyd1 treatment compared to the other three treatments (Fig. 4). The time course of expression levels reported in FlyBase for these genes ^56^, which is based on *Wolbachia*-infected flies ^57^, indicates that early embryos have lower levels than later embryo stages (see Table 4). Based on this pattern, a plausible hypothesis is that the Hyd1 treatment exhibits a developmental delay. To address this, we compared the expression levels among treatments of six genes that should exhibit high levels at the 0–2h embryo and substantially lower levels at the 2–4h embryo (see Table 4). Our rationale was that if the Hyd1 treatment were developmentally delayed, we might detect a trend of higher expression levels at these genes compared to the other treatments. Nonetheless, Hyd1 exhibited no such a pattern; except for one of these genes (i.e., FBgn0035955; not significant; sFig. 7). Therefore, collectively, the patterns of expression of Hyd1-infected embryos are inconsistent with a developmental delay.

**Table 4.** RPKM counts reported in FlyBase for the five DE genes found in our study and for six additional genes, “Developmental Markers” whose expression is higher in the 0-2h embryo than in later stages. Gene name or Annotation Symbol in parenthesis.

|  |  | FlyBase report for wMel infected flies (RPKM) |  |  |  |
| --- | --- | --- | --- | --- | --- |
|  |  | 0-2h embryo | 2-4h embryo | 4-6h embryo | 6-8h embryo |
| DE Genes in this study | FBgn0038339 (CG6118) | 1 | 1 | 1 | 3 |
|  | FBgn0000451 (*ect*) | 1 | 1 | 1 | 5 |
|  | FBgn0027527 (*Osi6*) | 10 | 7 | 8 | 31 |
|  | FBgn0037414 (*Osi6*) | 5 | 3 | 4 | 16 |
|  | FBgn0037099 (CG7173) | 0 | 0 | 1 | 2 |
| “Developmental markers” (high 0-2 h; much lower after) | FBgn0024991 (CG2694) | 56 | 9 | 4 | 7 |
|  | FBgn0035969 (CG4476) | 85 | 2 | 1 | 1 |
|  | FBgn0035955 (CG5194) | 56 | 2 | 1 | 1 |
|  | FBgn0038316 (CG6276) | 55 | 9 | 2 | 2 |
|  | FBgn0037807 (CG6293) | 94 | 7 | 1 | 1 |
|  | FBgn0030817 (CG4991) | 156 | 4 | 1 | 1 |

\* technical replicate

**Figure 7.**
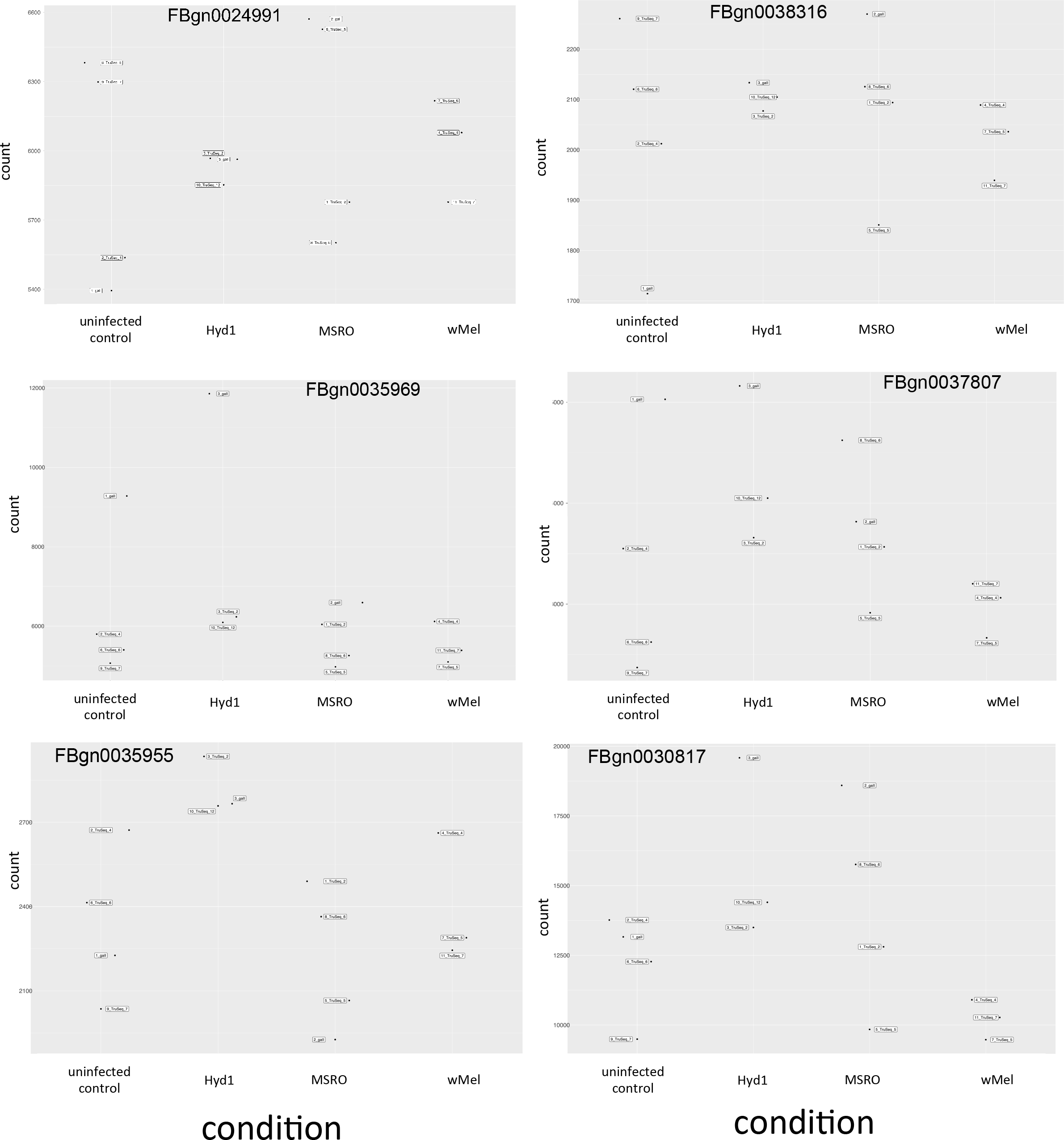
Normalized counts for each treatment and replicate, as estimated by Deseq2, for the six genes regarded as “Developmental markers” because they have high expression in the 0-2 h stage and much lower expression at subsequent stages. FlyBase-reported expression levels (in RPKM) are given in Table 4.

Available information on the function of the five genes at which the Hyd1 treatment had significantly lower expression provides little insight into the possible causes or consequences. Based on FlyBase, *ectodermal* (*ect*; FBgn0000451) is expected to have very low expression (RPKM = 1) at the stage we examined. It is expressed later (Stages 13–16 ≈ embryo 14-16h) in several tissue types: foregut, epidermis, trachea, and hindgut, with possible roles in cuticle development and tubular formation (e.g. tracheal tubes) ^58,59^. *Osiris 6* (*Osi6*; FBgn0027527) and *Osiris 7* (*Osi7*; FBgn0037414) are also expected to have very low expression (RPKM = 10 and 5, respectively) at the stage examined; with the highest expression at later embryonic stages (i.e., 14-16h). Both genes are of unknown function, but are physically close and belong to the Osiris gene cluster, which is present in insects and encodes a family of conserved putative transmembrane proteins of unknown function. In *D. melanogaster*, these genes are within the dosage-sensitive *Triploid-lethal* (*Tpl*) locus; individuals with one or three copies of *Tpl* die as late embryos or early first instar larvae ^60,61^. FBgn0038339 (CG6118) contains a BTB/POZ domain, and appears to be involved in regulation of transcription by RNA polymerase II. FBgn0037099 (CG7173) contains a Kazal domain (i.e., a type of serine proteinase inhibitor). Both of these genes are expected to have very low expression (RPKM = 1 and 0, respectively) at the stage we examined, with peak expression levels occurring in the 14-16h embryo.

Assuming that the significant results for the five genes are repeatable, they imply more perturbation of gene expression by Hyd1 than by the other symbiont strains; a finding that could reflect that *D. melanogaster* is outside the fundamental niche of Hyd1, as also suggested by its lower vertical transmission efficiency in this host (80%^62^), and by the difficulty of maintaining this artificial association in the lab (personal observation). A microarray-based expression analysis of adult *D. melanogaster* found that flies infected with Hyd1 in general exhibited less perturbation of gene expression than those infected by MSRO-Br and *Spiroplasma poulsonii* strain NSRO ^62^. The apparently different patterns of gene expression perturbation by Hyd1 of the two studies could be due to the different fly stages examined, different experimental tools, or other factors.

The lack of an effect of *w*Mel, a CI-inducing strain, on *D. melanogaster* early embryo mRNA composition, is not unexpected given what is known about the CI mechanism. Essentially, *w*Mel encodes two contiguous protein coding genes (*cifA* and *cifB*) ^18^. Each of these proteins is capable of causing CI when expressed in the male germline, whereas only cifA (when expressed in embryos) is able to rescue CI ^19^. Our results suggest that *w*Mel does not “prime” the egg for rescue by manipulating maternally-loaded transcripts or their degradation.

### 4.2 Depurination patterns

Our study capitalized on the presence of (non-target) rRNA reads in RNAseq datasets prepared by poly-A-tail enrichment, to estimate levels of depurination at the RIP target position of the sarcin-ricin loop (SLR) in the 28S rRNA of eukaryotes. We acknowledge that the levels of depurination estimated from mRNAseq data may be downwardly biased because rRNA depurinated by RIPs is highly prone to hydrolysis of the sugar-phosphate backbone at the lesion site ^63^. Furthermore, the freezing to which these samples were subjected between collection and library preparation might have decreased the detectability of depurination ^53^. In addition, it is not known whether the polyA-RNA enrichment protocol used prior to library preparation, which is aimed at depleting rRNA in the sample, could bias representation of depurinated vs. intact rRNA. Only one study has compared inferences of ribosome depurination from mRNAseq vs. qPCR assays in the same system. Based on mRNAseq, Hamilton et al. ^17^ detected ~3.8% depurination (i.e., ~96.2% adenine) in the nematode *Howardula* infesting adult *Drosophila neotestacea* harbouring the non-male-killing *Spiroplasma s*Neo strain. Comparatively, using the qPCR approach, the abundance of depurinated template representing RIP-induced depurination increased ~20-fold, whereas the levels of intact nematode rRNA were reduced ~six-fold in the presence of *Spiroplasma* ^17^. Notwithstanding the potential biases, the consistent finding of no depurination in *Spiroplasma*-free treatments vs. depurination in *Spiroplasma*-present treatments across three independent studies, lends credibility to the mRNAseq-based approach we employed for inference of depurination.

The significantly lower proportion of adenines at the RIP target site for the *Spiroplasma* treatments vs. the *Wolbachia* treatment and the control is consistent with the action of a *Spiroplasma*-encoded RIP, and supports recent findings that: (1) MSRO-UG infection causes ribosome depurination in *D. melanogaster* embryos; (2) that heterologous expression of a *Spiroplasma* RIP1 and RIP2 genes (separately) also causes ribosome depurination in *D. melanogaster* embryos (measured by a qPCR approach); and (3) that the degree of ribosome depurination (“RIP activity” *sensu* Garcia-Arraez, et al. ^38^) in the different *Spiroplasma* treatments or RIP transgene constructs is positively correlated with embryo mortality ^38^. Importantly, despite detecting significantly higher levels of depurinated template with the qPCR approach, none of the qPCR assays revealed significantly lower levels of intact template ^38^. This suggests that detectable depletion of intact template in the qPCR approach is not a pre-requisite for detecting a phenotypic effect of depurination.

Our results suggest that MSRO substrain Brazil (MSRO-BR) causes comparable levels of depurination to the Uganda substrain, which is consistent with its strong male-killing effect ^43,64^, and with the identical content of RIP-encoding genes (unpublished data). Interestingly, the male-killer MSRO generally exhibited a higher signal of depurination than the non-male-killer Hyd1 (MSRO range >1–8%; Hyd1 range <1–<3%; Fig. 6A). This difference could be the result of differences in titers of MSRO vs. Hyd1 at the embryonic stage. Densities of these strains in *D. melanogaster* Canton-S background at the adult stage are lower for Hyd1 than for MSRO-Br, but only immediately after adult eclosion ^64^; densities at the embryonic stage have not been compared. Alternatively, it is possible that the RIP genes encoded in the MSRO genome are more actively expressed or secreted, or are more efficient than the putative RIP genes detected in the genome of Hyd1 ^39^. Levels of fly ribosome depurination in the presence of *s*Neo, the non male-killing strain native to *D. neotestacea*, have not been assayed in embryos, but a qPCR assay of *D. neotestacea* ovaries indicated a small, albeit non significant, signal of depurination ^17^. Similarly, an mRNA-seq experiment of adult *D. neotestacea* revealed 0.4% depurination (i.e., %A = 99.6%) in the presence of *s*Neo ^17^. Therefore, the three members of the *poulsonii* clade examined to date (MSRO, Hyd1, and *s*Neo) encode RIPs capable of depurination of *Drosophila* ribosomes, but the male-killing strain exhibits the highest levels depurination, a phenomenon that appears to contribute to the male-killing mechanism ^38^. The general patterns, however, indicate that *Spiroplasma* RIPs are particularly efficient at depurinating the ribosomes of natural enemies of *Drosophila* (i.e., parasitic wasps and nematodes), leading to the hypothesis that RIP-induced depurination plays an important role in defence mechanism ^17,37^. The higher depurination levels of fly ribosomes detected in embryos (and ovaries) compared to other fly stages could be due to greater exposure of ribosomes to RIP prior to cellularization, after which *Spiroplasma* becomes effectively extra-cellular ^38^. The relatively high levels of depurination in old adults appears to be the result of higher *Spiroplasma* densities at that stage ^38^. Additional roles have been attributed to RIPs, which could contribute to the male-killing phenotype. For example, several RIPs are reported to cause DNA damage ^65,66^. Thus, one or more *Spiroplasma*-encoded RIP might directly contribute to the DNA damage reported during the process of male-killing ^30^.

## 5. Conclusions

This study employed a transcriptomics approach to examine whether cytoplasmically-transmitted bacteria, including reproductive manipulators that strongly impact survival of the embryonic stage of *Drosophila*, influence composition maternally-loaded mRNAs. The results revealed that mRNA composition does not differ significantly among the embryos harbouring the reproductive parasites *Spiroplasma* MSRO and *Wolbachia w*Mel and those lacking endosymbionts. Only the symbiont *Spiroplasma* Hyd1, which does not manipulate reproduction, appeared to alter expression levels of a handful of genes (5–12), but not all analytical approaches supported this finding. Our power analyses indicated that our experimental design should be able to detect most of the genes exhibiting ≥ 3-log-fold change in expression among treatments. Collectively, our results suggest that these cytoplasmically-transmitted bacteria do not alter the composition of mRNAs of the early embryo, and are thus unlikely to use this mechanism to exert their reproductive phenotypes. Capitalizing on several trascriptomics datasets, this study also detected signals of *Spiroplasma*-induced damage to ribosomes in early *Drosophila* embryos, with greater damage caused by the male-killer (MSRO) than the non-male-killer (Hyd1), consistent with recent results implicating this mechanism in male killing.

## Supporting information

Description of Supplemental Files

## Acknowledgements

Library preparation and Illumina sequencing were performed by the Texas AgriLife Genomics and Bioinformatics facility. Portions of this research were conducted with high performance research computing resources provided by Texas A&M University (https://hprc.tamu.edu) and UNAM-CCG (litza server). This work was supported by National Institutes of Health grant R03 AI078348 to MM. Lacie L. Güenther and Jialei Xie provided technical assistance.

## Author contributions

Conceived and designed study: NOS, MM, RA, JWE

Conducted the experiments: NOS, MM

Analysed the data: MM, NOS, PR, VMHA, RA

Interpreted results: MM, NOS, PR, VMHA, RA, JWE

Drafted the manuscript: MM, NOS, PR, VMHA

Reviewed and edited the manuscript: MM, NOS, PR, VMHA, RA, JWE

## Data availability statement

The datasets generated during and/or analysed during the current study are available as Supporting Data or at NCBI’s Sequence Read Archive (SRA) database under Accession Numbers SRR7279355–SRR7279369 (BioProject PRJNA474708; BioSample SAMN09370647).

