## Supplementary material for "Effect of heritable symbionts on maternally-derived embryo transcripts": Description of Supplemental Files

**Description of Supplementary Information Files**

**Supporting Table S1.** Number of reads that mapped to the site within the sarcin-ricin-loop (SRL) of the 28S rRNA gene that is targeted by ribosome inactivating proteins (RIPs), and number of reads that contained an adenine (expected in the absence of RIP activity) or other base (expected as a result of depurination) at the target position.

**Supporting Table S2**. Number and proportion of reads in symbiont treatments that mapped to the corresponding symbiont genome. Numbers and proportion of reads that mapped to the ribosomal RNA genes of the symbionts.

**Supporting Table S3**. Metadata of reads submitted to NCBI including BioProject and BioSample numbers.

**Supporting Data File S1.** Compressed archive containing the reference genome (dmel-all-r6.18) and the command lines used to run the depurination analyses and each of the differential expression analyses pipelines. Files included: ReadMapping_Counting.txt; edgeR_exon.R; limma_exon.R; deseq_exon.R; deseq_gene.R; limma_gene.R; edgeR_gene.R; dmel-all-r6.18.gff.gz

**Supporting Data File S2.** Compressed archive containing the Htseq-generated raw counts for each gene (file: 2drosophila-htseq_counts_gene-1.txt) and exon (file: exon-drosophila-htseq_counts_all-1.txt).

**Supporting Data File S3.** Compressed archive containing sample definition files used for the edgeR and limma analyses (SampleInfo.txt) and for the DeSeq analyses (2SampleInfo.txt).

**Supporting Data File S4.** Compressed archive containing the main output file from each of the differential expression analyses. Files included are:

gene_edgeR_results.xlsx: contains the edgeR results based on genes using glm and glmqlf

gene_deseq2_results.xlsx: contains the deseq2 results based on genes.

gene_limma_voom_results.xlsx: contains the Limma-Voom results based on genes.

exon_limma_results.xlsx: contains the Limma-Voom results based on genes.

exon_Deseq2_results.xlsx: contains the deseq2 results based on exons.

exon_edgeR_n_4_results.xlsx: contains the edgeR results of glm and glmqlf for analyses based on exons.
