## Supporting Data File S1 for "Effect of heritable symbionts on maternally-derived embryo transcripts": deseq_exon.R

```
directory <- "/Users/paulinoramirez/R_program/RNA-seq/exon/htseq"
sampleFiles <- grep("*.txt",list.files(directory),value=TRUE)
sampleCondition <- read.delim("2SampleInfo.txt")
sampleTable <- data.frame(sampleName = sampleCondition$X,
                          fileName = sampleCondition$Filename,
                          condition = sampleCondition$condition)

library("DESeq2")
ddsHTSeq <- DESeqDataSetFromHTSeqCount(sampleTable = sampleTable,
                                       directory = directory,
                                       design= ~ condition)
ddsHTSeq

#pre-filter low count genes
keep <- rowSums(counts(ddsHTSeq)) >= 10
ddsHTSeq <- ddsHTSeq[keep,]

#specifying the reference level:
ddsHTSeq$condition <- relevel(ddsHTSeq$condition, ref = "U")

ddsHTSeq <- DESeq(ddsHTSeq)
res <- results(ddsHTSeq)
res
###
res_MSRO <- results(ddsHTSeq, contrast=c("condition","MSRO","U"))
resOrdered_MSRO <- res_MSRO[order(res_MSRO$pvalue),];resOrdered_MSRO
summary(res_MSRO)


write.csv(as.data.frame(resOrdered_MSRO), 
          file="condition_MSRO_results-exon.csv")

res_hy1 <- results(ddsHTSeq, contrast=c("condition","hy1","U")) 
resOrdered_hy1 <- res_hy1[order(res_hy1$pvalue),];resOrdered_hy1
summary(res_hy1)


res_wmel <- results(ddsHTSeq, contrast=c("condition","wmel","U"))
resOrdered_wmel <- res_wmel[order(res_wmel$pvalue),];resOrdered_wmel
summary(res_wmel)


write.csv(as.data.frame(resOrdered_hy1), 
          file="condition_hy1_results-exon.csv")
write.csv(as.data.frame(resOrdered_wmel), 
          file="condition_wmel_results-exon.csv")


resLFC_hy1 <- lfcShrink(ddsHTSeq, coef="condition_hy1_vs_U");resLFC_hy1
resApe_hy1 <- lfcShrink(ddsHTSeq, coef="condition_hy1_vs_U", type="apeglm")
resAsh_hy1 <- lfcShrink(ddsHTSeq, coef="condition_hy1_vs_U", type="ashr")

par(mfrow=c(1,3), mar=c(4,4,2,1))
xlim <- c(1,5); ylim <- c(-1,1)
plotMA(resLFC_hy1, xlim=xlim, ylim=ylim, main="normal")
plotMA(resApe_hy1, xlim=xlim, ylim=ylim, main="apeglm")
plotMA(resAsh_hy1, xlim=xlim, ylim=ylim, main="ashr")
par(mfrow=c(1,1))


plotCounts(ddsHTSeq, gene="FBgn0038339", intgroup="condition")
plotCounts(ddsHTSeq, gene="FBgn0000451", intgroup="condition")
plotCounts(ddsHTSeq, gene="FBgn0027527", intgroup="condition")
plotCounts(ddsHTSeq, gene="FBgn0037414", intgroup="condition")
plotCounts(ddsHTSeq, gene="FBgn0001254", intgroup="condition")

resSig_hy1 <- subset(resOrdered_hy1, padj < 0.1)
resSig_hy1
write.csv(as.data.frame(resSig_hy1), 
          file="exon_condition_hy1_results.csv")
```
