## Supporting Data File S1 for "Effect of heritable symbionts on maternally-derived embryo transcripts": deseq_gene.R

```
setwd("~/R_program/RNA-seq/gene/deseq2")
directory <- "/Users/paulinoramirez/R_program/RNA-seq/gene/htseq_counts"
sampleFiles <- grep("*.txt",list.files(directory),value=TRUE)
sampleCondition <- read.delim("2SampleInfo.txt")
sampleTable <- data.frame(sampleName = sampleCondition$X,
                          fileName = sampleCondition$Filename,
                          condition = sampleCondition$condition)

library("DESeq2")
ddsHTSeq <- DESeqDataSetFromHTSeqCount(sampleTable = sampleTable,
                                       directory = directory,
                                       design= ~ condition)
ddsHTSeq

#pre-filter low count genes
keep <- rowSums(counts(ddsHTSeq)) >= 10
ddsHTSeq <- ddsHTSeq[keep,]

#specifying the reference level:
ddsHTSeq$condition <- relevel(ddsHTSeq$condition, ref = "U")

ddsHTSeq <- DESeq(ddsHTSeq)
res <- results(ddsHTSeq)
res
###
res_MSRO <- results(ddsHTSeq, contrast=c("condition","MSRO","U"))
resOrdered_MSRO <- res_MSRO[order(res_MSRO$pvalue),];resOrdered_MSRO
summary(res_MSRO)
sum(res_MSRO$padj < 0.1, na.rm=TRUE)

write.csv(as.data.frame(resOrdered_MSRO), 
          file="condition_MSRO_results.csv")

res_hy1 <- results(ddsHTSeq, contrast=c("condition","hy1","U")) 
resOrdered_hy1 <- res_hy1[order(res_hy1$pvalue),];resOrdered_hy1


res_wmel <- results(ddsHTSeq, contrast=c("condition","wmel","U"))
resOrdered_wmel <- res_wmel[order(res_wmel$pvalue),];resOrdered_wmel


write.csv(as.data.frame(resOrdered_hy1), 
          file="condition_hy1_results.csv")
write.csv(as.data.frame(resOrdered_wmel), 
          file="condition_wmel_results.csv")

#####Visualization
resultsNames(ddsHTSeq)

resLFC_hy1 <- lfcShrink(ddsHTSeq, coef="condition_hy1_vs_U");resLFC_hy1
resApe_hy1 <- lfcShrink(ddsHTSeq, coef="condition_hy1_vs_U", type="apeglm")
resAsh_hy1 <- lfcShrink(ddsHTSeq, coef="condition_hy1_vs_U", type="ashr")
plotMA(resLFC_hy1, ylim=c(-2,2))

par(mfrow=c(1,3), mar=c(4,4,2,1))
xlim <- c(1,5); ylim <- c(-3,3)
plotMA(resLFC_hy1, xlim=xlim, ylim=ylim, main="normal")
plotMA(resApe_hy1, xlim=xlim, ylim=ylim, main="apeglm")
plotMA(resAsh_hy1, xlim=xlim, ylim=ylim, main="ashr")
par(mfrow=c(1,1))


plotCounts(ddsHTSeq, gene="FBgn0038339", intgroup="condition")
plotCounts(ddsHTSeq, gene="FBgn0000451", intgroup="condition")
plotCounts(ddsHTSeq, gene="FBgn0027527", intgroup="condition")
plotCounts(ddsHTSeq, gene="FBgn0037414", intgroup="condition")
plotCounts(ddsHTSeq, gene="FBgn0037099", intgroup="condition")
plotCounts(ddsHTSeq, gene="FBgn0001254", intgroup="condition")
plotCounts(ddsHTSeq, gene="FBgn0000568", intgroup="condition")
plotCounts(ddsHTSeq, gene="FBgn0001256", intgroup="condition")
plotCounts(ddsHTSeq, gene="FBgn0031505", intgroup="condition")


resSig_hy1 <- subset(resOrdered_hy1, padj < 0.1)
resSig_hy1
write.csv(as.data.frame(resSig_hy1), 
          file="condition_hy1_results.csv")

#################

vsd <- vst(ddsHTSeq, blind=FALSE)
rld <- rlog(ddsHTSeq, blind=FALSE)
head(assay(vsd), 3)


##########
# this gives log2(n + 1)
ntd <- normTransform(ddsHTSeq)
library("vsn")
meanSdPlot(assay(ntd))
meanSdPlot(assay(vsd))
meanSdPlot(assay(rld))
# 
library("pheatmap")
select <- order(rowMeans(counts(ddsHTSeq,normalized=TRUE)),decreasing=TRUE)[1:20]
df <- as.data.frame(colData(ddsHTSeq))

rownames(df) 
rownames(df) <- df[,1]

pheatmap(assay(ntd)[select,], cluster_rows=FALSE, show_rownames=FALSE,cluster_cols=FALSE, annotation_col=df)
pheatmap(assay(vsd)[select,], cluster_rows=FALSE, show_rownames=FALSE,cluster_cols=FALSE, annotation_col=df)
pheatmap(assay(rld)[select,], cluster_rows=FALSE, show_rownames=FALSE,cluster_cols=FALSE, annotation_col=df)

sampleDists <- dist(t(assay(vsd)))
library("RColorBrewer")
sampleDistMatrix <- as.matrix(sampleDists)
rownames(sampleDistMatrix) <- paste(vsd$condition, vsd$type, sep="-")
colnames(sampleDistMatrix) <- NULL
colors <- colorRampPalette( rev(brewer.pal(9, "Blues")) )(255)
pheatmap(sampleDistMatrix,
         clustering_distance_rows=sampleDists,
         clustering_distance_cols=sampleDists,
         col=colors)


plotDispEsts(ddsHTSeq)

par(mar=c(8,5,2,2))
boxplot(log10(assays(ddsHTSeq)[["cooks"]]), range=0, las=2)


plotPCA(vsd, intgroup=c("condition"))
z <- plotPCA(vsd, intgroup=c("condition"))
z + geom_label(aes(label = name))


sessionInfo()

plotCounts(ddsHTSeq, gene="FBgn0263352", intgroup="condition")
plotCounts(ddsHTSeq, gene="FBgn0261068", intgroup="condition")
plotCounts(ddsHTSeq, gene="FBgn0004650", intgroup="condition")
plotCounts(ddsHTSeq, gene="FBgn0036741", intgroup="condition")
plotCounts(ddsHTSeq, gene="FBgn0031505", intgroup="condition")
plotCounts(ddsHTSeq, gene="FBgn0025839", intgroup="condition")
plotCounts(ddsHTSeq, gene="FBgn0262601", intgroup="condition")
plotCounts(ddsHTSeq, gene="FBgn0015393", intgroup="condition")
plotCounts(ddsHTSeq, gene="FBgn0062928", intgroup="condition")
plotCounts(ddsHTSeq, gene="FBgn0063670", intgroup="condition")
plotCounts(ddsHTSeq, gene="FBgn0032242", intgroup="condition")
plotCounts(ddsHTSeq, gene="FBgn0034240", intgroup="condition")
plotCounts(ddsHTSeq, gene="FBgn0031263", intgroup="condition")
plotCounts(ddsHTSeq, gene="FBgn0031677", intgroup="condition")


goodHosp <- c("FBgn0263352",
"FBgn0261068",
"FBgn0004650",
"FBgn0036741",
"FBgn0031505",
"FBgn0025839",
"FBgn0262601",
"FBgn0015393",
"FBgn0062928",
"FBgn0063670",
"FBgn0032242",
"FBgn0034240",
"FBgn0031263",
"FBgn0031677",
"FBgn0040007",
"FBgn0002579",
"FBgn0010078",
"FBgn0032518",
"FBgn0003274",
"FBgn0044511",
"FBgn0031231")

x = counts_deseq[goodHosp,]         


counts_deseq = counts(ddsHTSeq,normalized = TRUE)


write.csv(as.data.frame(x), 
          file="counts_deseq.csv")
```
