## Supporting Data File S1 for "Effect of heritable symbionts on maternally-derived embryo transcripts": edgeR_exon.R

```
library(edgeR)
library(limma)
library(Glimma)
library(gplots)
library(org.Mm.eg.db)
library(RColorBrewer)
library(splines)


Counts = read.table("exon-drosophila-htseq_counts_all-1.txt", header=T)
head(Counts)
dim(Counts)
# Read the sample information into R
sampleinfo <- read.delim("SampleInfo.txt")


##most expressed gene is FBgn0013686(16sRNA) and FBgn0284245(ef-1α) 
##If treated as outliers and removed
Counts <- Counts[-match("FBgn0013686", rownames(Counts)),]
Counts <- Counts[-match("FBgn0284245", rownames(Counts)),]


##Creating a DGEList objec
dgList <- DGEList(counts=Counts, genes=rownames(Counts))
dgList$samples
head(dgList$counts)
head(dgList$genes)

#Filtering cpm less than  0.5 
countsPerMillion <- cpm(dgList)
summary(countsPerMillion)
countCheck <- countsPerMillion > 0.5
head(countCheck)
##### sample replicates = 3
keep <- which(rowSums(countCheck) >= 4)
dgList <- dgList[keep,]
summary(cpm(dgList))


#Normalisation by the TMM (method) EdgeR Pipeline
dgList <- calcNormFactors(dgList, method="TMM")

## Graph building
col.cell <- c("purple","orange","blue","red")[sampleinfo$Type]
data.frame(sampleinfo$Type,col.cell)
#Data Exploration
plotMDS(dgList,col=col.cell)
legend("topleft",fill=c("purple","orange","blue","red"),legend=levels(sampleinfo$Type))
title("MDS-Gene,N=3,CPM > 0.5")

#### make a Count matrix
group <- (sampleinfo$Type)
designMat <- model.matrix(~ 0 + group, data=sampleinfo$Type)
designMat


### Normalize data generatated from TMM
y <- dgList
y <- calcNormFactors(y)
y$samples


# qCML method is only applicable on datasets with a single factor design since it fails to take into account 
#the effects from multiple factors in a more complicated experiment

# estimate common dispersion and tagwise dispersions in one run 
y <- estimateDisp(y,designMat)
#glmQLFit - quasi likilyhood
fit_glmQl <- glmQLFit(y, designMat)
#general linearized model
fit_glm <- glmFit(y, designMat)

#######MSR0VsControll - Only output first top 20 de genes
#quasi likelihood 
qlf_MSRO <- glmQLFTest(fit_glmQl, contrast=c(0,1,-1,0))
topTags(qlf_MSRO,n=Inf)
MSROvsUnifected=topTags(qlf_MSRO,n=Inf)
write.table(MSROvsUnifected,file="exon-MSROvsUnifected-glmqlf-gene_n_4_April_2018.tsv",sep="\t",quote=F)

#GLM_model
lrt_MSRO <- glmLRT(fit_glm,contrast=c(0,1,-1,0))
topTags(lrt_MSRO,n=Inf)
MSROvsUninfected=topTags(lrt_MSRO,n=Inf)
write.table(MSROvsUninfected,file="exon-MSROvsUninfected-glm-gene_n_4_April_2018.tsv",sep="\t",quote=F)


####### hyd1VsControll

### QLFT test
qlf_hyd1 <- glmQLFTest(fit_glmQl, contrast=c(1,0,-1,0))
topTags(qlf_hyd1,n=Inf)
hyd1vsUninfected=topTags(qlf_hyd1,n=Inf)
write.table(hyd1vsUninfected,file="exon-hyd1vsUninfected-glmqlf-gene_n_4_April_2018.tsv",sep="\t",quote=F)


### LRT test 
lrt_hyd1 <- glmLRT(fit_glm,contrast=c(1,0,-1,0))
topTags(lrt_hyd1,n=Inf)
hyd1vsUninfected=topTags(lrt_hyd1,n=Inf)
write.table(hyd1vsUninfected,file="exon-hyd1vsUninfected-glm-gene_n_4_April_2018.tsv",sep="\t",quote=F)

#Plot biological coeffcient of variation
plotBCV(y)

plotQLDisp(fit_glmQl)


#######wMelVsControll
 qlf_wol <- glmQLFTest(fit_glmQl, contrast=c(0,0,-1,1))
 topTags(qlf_wol,n=Inf)
 wMelvsUnifected=topTags(qlf_wol,n=Inf)
 write.table(wMelvsUnifected,file="exon-wMelvsUnifected-glmqlf-gene_n_4_April_2018.tsv",sep="\t",quote=F)
 
 lrt_wol <- glmLRT(fit_glm,contrast=c(0,0,-1,1))
 topTags(lrt_wol,n=Inf)
 hyd1vsUninfected=topTags(lrt_wol,n=Inf)
 write.table(hyd1vsUninfected,file="exon-wmelvsUninfected-glm-gene_n_4_April_2018.tsv",sep="\t",quote=F)
########Anova-one ways

design <- model.matrix(~group,data=sampleinfo$Type)
design


qlf_a <- glmQLFTest(fit_glmQl, coef=1:4)
topTags(qlf_a, n=50)
anova_all=topTags(qlf_a,n=500)
write.table(anova_all,file="anova-all-DE_gene_n_4_April.tsv",sep="\t",quote=F)


####Output Version Info and packages used
sessionInfo()
```
