## Supporting Data File S1 for "Effect of heritable symbionts on maternally-derived embryo transcripts": limma_exon.R

```
library(edgeR)
library(limma)
library(Glimma)
library(gplots)
library(org.Mm.eg.db)
library(RColorBrewer)

# Read the data into R
seqdata <- read.delim("exon-drosophila-htseq_counts_all-1.txt", stringsAsFactors = FALSE)
# Read the sample information into R
sampleinfo <- read.delim("SampleInfo.txt")
sampleinfo

# Remove first  columnsfrom seqdata
countdata <- seqdata[,-(1)]
# Look at the output
head(countdata)

# Store GeneID as rownames
rownames(countdata) <- seqdata[,1]

##most expressed gene is FBgn0013686(16sRNA) and FBgn0284245(ef-1α) 
##If treated as outliers and removed
countdata <- countdata[-match("FBgn0013686", rownames(countdata)),]
countdata <- countdata[-match("FBgn0284245", rownames(countdata)),]


#Now take a look at the column names
colnames(countdata)

# using substr, you extract the characters starting at position 1 and stopping at position 7 of the colnames
#colnames(countdata) <- substr(colnames(countdata),start=1,stop=7)
head(countdata)


table(colnames(countdata)==sampleinfo$Filename)


#We’ll use the cpm function from the edgeR library (M D Robinson, McCarthy, and Smyth 2010) to generate the CPM values and then filter. Note that by converting to CPMs we are normalising for the different sequencing depths for each sample.
#Obtain CPMs
myCPM <- cpm(countdata)
# Have a look at the output
head(myCPM)

# Which values in myCPM are greater than 0.5 
thresh <- myCPM > 0.5

# This produces a logical matrix with TRUEs and FALSEs
head(thresh)

# Summary of how many TRUEs there are in each row
table(rowSums(thresh))

##When there are biological replicates in each group, 
##in this case we have a sample size of 4 in each group, 
##we favour filtering on a minimum counts per million threshold present in at least 4 samples

## we would like to keep genes that have at least 4 TRUES in each row of thresh
keep <- rowSums(thresh) >= 4
## Subset the rows of countdata to keep the more highly expressed genes
counts.keep <- countdata[keep,]
summary(keep)


dim(counts.keep)


## Let's have a look and see whether our threshold of 0.5 does indeed correspond to a count of about 10-15
## We will look at the first sample
plot(myCPM[,1],countdata[,1])

## Let us limit the x and y-axis so we can actually look to see what is happening at the smaller counts
plot(myCPM[,1],countdata[,1],ylim=c(0,200),xlim=c(0,3))
## Add a vertical line at 0.5 CPM
abline(v=3.0)
abline(h=100)


##Next we’ll create a DGEList object. This is an object used by edgeR to store count data. It has a number of slots for storing various parameters about the data.

y <- DGEList(counts.keep)
## have a look at y
y

## See what slots are stored in y
names(y)


## Library size information is stored in the samples slot
y$samples


## Library size information is stored in the samples slot
y$samples$lib.size

### Now that we have got rid of the lowly expressed genes and have our counts stored in a DGEList object, we can look at a few different plots to check that the data is good quality, and that the samples are as we would expect.
## The names argument tells the barplot to use the sample names on the x-axis
## The las argument rotates the axis names
barplot(y$samples$lib.size,names=colnames(y),las=2)
# Add a title to the plot
title("Barplot of Raw library sizes (Gene)")

# Get log2 counts per million
logcounts <- cpm(y,log=TRUE)
# Check distributions of samples using boxplots
boxplot(logcounts, xlab="", ylab="Log2 counts per million",las=2)
# Let's add a blue horizontal line that corresponds to the median logCPM
abline(h=median(logcounts),col="blue")
title("Boxplots of logCPMs- N=4")

###Multidimensional scaling plots (Batch effects?)
levels(sampleinfo$Type)
## Let's choose purple for basal and orange for luminal
col.cell <- c("purple","orange","blue","red")[sampleinfo$Type]
data.frame(sampleinfo$Type,col.cell)
plotMDS(y,col=col.cell)
legend("topleft",fill=c("purple","orange","blue","red"),legend=levels(sampleinfo$Type))
# Add a title
title("Infection Type-gene N=4")

###Hierarchical clustering with heatmaps
## We estimate the variance for each row in the logcounts matrix
var_genes <- apply(logcounts, 1, var)
head(var_genes)

## Get the gene names for the top 100 most variable genes
select_var <- names(sort(var_genes, decreasing=TRUE))[1:30]
head(select_var)

## Subset logcounts matrix
highly_variable_lcpm <- logcounts[select_var,]
dim(highly_variable_lcpm)
head(highly_variable_lcpm)

## Get some nicer colours
mypalette <- brewer.pal(11,"RdYlBu")
morecols <- colorRampPalette(mypalette)
# Set up colour vector for type variable
col.cell <- c("purple","orange","blue","red")[sampleinfo$Type]

# Plot the heatmap
heatmap.2(highly_variable_lcpm,col=rev(morecols(50)),trace="none", main="Top 100 most variable genes across ",ColSideColors=col.cell,scale="row")
# Save the heatmap


###Normalisation for composition bias

## Apply normalisation to DGEList object
y <- calcNormFactors(y)
##This will update the normalisation factors in the DGEList object (their default values are 1). Take a look at the normalisation factors for these samples

y$samples

## composition bias problem
par(mfrow=c(1,2))
plotMD(logcounts,column = 9)
abline(h=0,col="grey")
plotMD(logcounts,column = 2)
abline(h=0,col="grey")

#The mean-difference plots show average expression (mean: x-axis) against log-fold-changes (difference: y-axis). Because our DGEList object contains the normalisation factors,
#if we redo these plots using y, we should see the composition bias problem has been solved.

par(mfrow=c(1,2))
plotMD(logcounts,column = 2)
abline(h=0,col="grey")
plotMD(logcounts,column = 9)
abline(h=0,col="grey")
plotMD(y,column = 2)
abline(h=0,col="grey")
plotMD(y,column = 9)
abline(h=0,col="grey")

#####Repeat graphs with normalized y

n_y <- y

## Library size information is stored in the samples slot  NOTE NEW norm.factors
n_y$samples


## Library size information is stored in the samples slot
n_y$samples$lib.size


#######Differential expression with limma-voom
##make "group" variable
group <- (sampleinfo$Type)
## Look at group variable again
group

## Specify a design matrix without an intercept term
design <- model.matrix(~ 0 + group)
design 


## Make the column names of the design matrix a bit nicer
colnames(design) <- levels(group)
design


## Voom transform the data
## Once we have our design matrix ready to go, we can perform our voom transformation. 
## Voom will automatically adjust the library sizes using the norm.factors already calculated. 
## The voom transformation uses the experiment design matrix, and produces an EList object. 
## We can add plot=TRUE to generate a plot of the mean-variance trend. 
## This plot can also tell us if there are any genes that look really variable in our data, and if we’ve filtered the low counts adequately.
## 
##vroom
par(mfrow=c(1,1))
v <- voom(n_y,design,plot = TRUE)
## We can repeat the box plots for the normalised data to compare to before normalisation. 
## The expression values in v$E are already log2 values so we don’t need to log-transform.
par(mfrow=c(1,2))
boxplot(logcounts, xlab="", ylab="Log2 counts per million",las=2,main="Unnormalised logCPM")
## Let's add a blue horizontal line that corresponds to the median logCPM
abline(h=median(logcounts),col="blue")
boxplot(v$E, xlab="", ylab="Log2 counts per million",las=2,main="Voom transformed logCPM")
## Let's add a blue horizontal line that corresponds to the median logCPM
abline(h=median(v$E),col="blue")

## Testing for differential expression
## Now that we have the voom transformed data we can use limma to test for differential expression.
## First we fit a linear model for each gene using the lmFit function in limma. 
## lmFit needs the voom object and the design matrix that we have already specified, 
## which is stored within the voom object.
## Fit the linear model
fit <- lmFit(v)
names(fit)

## lmFit estimates group means according to the design matrix, as well as gene-wise variances. 
## Since we are interested in differences between groups, we need to specify which comparisons we want to test. 
## The comparison of interest can be specified using the makeContrasts function.
## This is done by defining the null hypothesis as MSRO - U = 0 for each gene.
## Note that the group names must exactly match the column names of the design matrix.

cont.matrix <- makeContrasts(MSROVscontroll=MSRO - U, hy1Vscontroll=hy1 - U, wmelVscontroll=wmel - U, levels=design)
cont.matrix

## Now we can apply the contrasts matrix to the fit object to get the statistics and estimated parameters of our comparison that we are interested in.
## Here we call the contrasts.fit function in limma.
fit.cont <- contrasts.fit(fit, cont.matrix)
##The final step is to call the eBayes function, which performs empirical Bayes shrinkage on the variances, and estimates moderated t-statistics and the associated p-values.
fit.cont <- eBayes(fit.cont)
##Check the dimensions of the fit object
dim(fit.cont)

##We can use the limma decideTests function to generate a quick summary of DE genes for the contrasts
summa.fit <- decideTests(fit.cont)
summary(summa.fit)

#hy1Vscontroll
x=topTable(fit.cont,coef="hy1Vscontroll",sort.by="p",number=Inf)
#MSROVscontroll
y=topTable(fit.cont,coef="MSROVscontroll",sort.by="p",number=Inf)
#wmel1Vscontroll
z=topTable(fit.cont,coef="wmelVscontroll",sort.by="p",number=Inf)

write.table(x,file="exon-hy1vsUnifected-limma.tsv",sep="\t",quote=F)
write.table(y,file="exon-MSROvsUnifected-limma.tsv",sep="\t",quote=F)
write.table(z,file="exon-wMelvsUnifected-limma.tsv",sep="\t",quote=F)


# We want to highlight the significant genes. We can get this from decideTests.
par(mfrow=c(1,1))
plotMD(fit.cont,coef=1,status=summa.fit[,"hy1Vscontroll"])

plotMD(fit.cont,coef=1,status=summa.fit[,"MSROVscontroll"])

plotMD(fit.cont,coef=1,status=summa.fit[,"wmelVscontroll"])


# For the volcano plot we have to specify how many of the top genes to hightlight.
# We can also specify that we want to plot the gene symbol for the highlighted genes.
# let's highlight the top 100 most DE genes
volcanoplot(fit.cont,coef=1,highlight=1,names=fit.cont$genes$SYMBOL)


sessionInfo()
```
