## Supporting Data File S1 for "Effect of heritable symbionts on maternally-derived embryo transcripts": ReadMapping_Counting.txt

######## mRNA seq code


######## read mapping
hisat2 -q -p $threads -x $prefix  -U $fastq_file  -S $Sam

######## gene or exon counts
htseq-count --stranded=no --type= <exon or gene > $alignment_file_1 $gff_file > ${prefix_1}_count_exon_table.txt


######## code for depurination analyses

######## read mapping to the 28S rRNA gene of Drosophila melanogaster

bowtie2 -q -p 10 -x index_28S -U file.fastq -S output.sam 

######## Code to extract mapped reads

samtools view -S -F4  File.sam -o  output.sam

######## code to convert sam to fastq format

File.sam | grep -v ^@ | awk '{print "@"$1"\n"$10"\n+\n"$11}' >  File.fastq
